# Higher bat and bird γ-diversity in structurally complex forests is driven by distinct α- and β-diversity responses

**DOI:** 10.1101/2025.08.25.671712

**Authors:** Clara Wild, Anne Chao, Po-Yen Chuang, Marc Cadotte, Nico Daume, Orsi Decker, Ronja Hausmann, Sophia Hochrein, Michael Junginger, Mareike Kortmann, Sonja Kümmet, Soumen Mallick, Oliver Mitesser, Ruth Pickert, Julia Rothacher, Kai Sattler, Jens Schlüter, Simon Thorn, Jörg Müller

## Abstract

Effective conservation management and habitat restoration rely on understanding how biodiversity responds to environmental change. Centuries of silviculture have homogenized forests and their species communities globally, reducing biodiversity. To test whether restoring forest structural complexity can promote biodiversity, we conducted a large-scale, spatially explicit landscape experiment. At 11 sites across Germany, we compared bat and bird diversity in forests with experimentally enhanced heterogeneity by increasing deadwood and canopy complexity to homogeneous production forests. Both taxa were investigated by autonomous acoustic recorders and automatic species identification. We quantified within-patch (α-), between-patch (β-), and landscape-level (γ-) diversity, emphasizing infrequent to highly frequent species for taxonomic, functional, and phylogenetic diversity. The pairwise comparisons of the sites were synthesized using a newly developed meta-analysis of rarefaction-extrapolation curves. γ-diversity increased significantly in structurally heterogeneous forests for both taxa, albeit through distinct taxon-specific mechanisms. Bat γ-diversity gains were primarily driven by higher β-diversity, indicating greater dissimilarity in species assemblages among patches, while bird γ-diversity increased via higher α-diversity within patches. Bat diversity increases were mainly taxonomic, suggesting functional similarity in the communities, whereas birds showed the highest gains in functional diversity, indicating that experimental treatments resulted in greater trait dissimilarity. Our results provide experimental evidence under real-world conditions that γ-diversity can be shaped by different diversity mechanisms. These patterns likely originate from differences in activity ranges, such as the large-scale movements of foraging bats in contrast to the more spatially restricted, territorial behavior of birds. This highlights the need for taxon-specific restoration strategies in homogenized landscapes.

## Introduction

Biodiversity is changing globally, often resulting in declines in ecosystem functioning and stability at the local scale, with serious consequences for both wildlife and human well-being (Sala et al. 2000, Cardinale et al. 2012, Gossner et al. 2016). Human pressures, especially intensive land use, have traditionally been associated with reduced local species diversity, shifts in community composition, and biotic homogenization (McKinney and Lockwood 1999, Olden and Rooney 2006, Hooper et al. 2012, Jaureguiberry et al. 2022). Despite decades of research, the impacts of anthropogenic drivers can be complex, making predictions of biodiversity patterns difficult. Recent studies have challenged long-held assumptions that human impacts lead to universal patterns of homogenization and landscape fragmentation effects, reinforcing that biodiversity responses are highly taxon-, habitat-, and scale-dependent (Gonçalves-Souza et al. 2025, Keck et al. 2025). Moreover, biodiversity is multi-faceted, with patterns varying among rare, common, and dominant species within species assemblages, captured by orders of q in Hill numbers (Chao et al. 2014, McGill et al. 2015). Understanding the different mechanisms driving biodiversity change across different dimensions of diversity and spatial scales is essential for developing effective restoration strategies, especially those aimed at increasing habitat heterogeneity.

Throughout large parts of Europe, centuries of economically oriented silviculture have produced structurally simplified, even-aged forest landscapes that lack both early and late successional stages. Yet, these two stages provide sunny conditions and deadwood build-up, respectively, and both are critical for supporting biodiversity (Brunet et al. 2010, Hilmers et al. 2018, Aszalós et al. 2022). The generality of a positive relationship between environmental heterogeneity, such as structural complexity, and diversity has been widely debated (MacArthur and MacArthur 1961, Lundholm 2009, Stein et al. 2014). According to the habitat heterogeneity hypothesis, structurally complex habitats support more species, up to a certain area threshold, by offering a greater variety of niches (MacArthur and MacArthur 1961, Allouche et al. 2012, Stein et al. 2014, Heidrich et al. 2020). While previous studies support this relationship for bats and birds, with strong evidence for vertical and particularly for horizontal structures (Renner et al. 2018, Heidrich et al. 2020, Rigo et al. 2024), empirical evidence that spans multiple scales, regions, and controlled experimental settings remains scarce. Addressing this challenge requires large-scale, nested study designs and analytical approaches, such as standardized meta-analyses, that ensure consistency across regions and account for differences in sample completeness, both of which we present in this study. Moreover, habitat heterogeneity can affect within-patch α- and between-patch β-diversity in contrasting ways, diverging across dissimilar habitats or aligning in more similar or connected ones, ultimately shaping landscape-level γ-diversity (Pastro et al. 2011). Dissimilarities between species assemblages can arise from different processes and are most commonly attributed to species turnover, that is, the replacement of species (Legendre 2014, Soininen et al. 2018). However, little is known about the relationship dynamics between α-, β-, and γ-diversity, and the role of α- and β-diversity in shaping γ in a controlled experiment.

Beyond species richness, understanding functional diversity is crucial for biodiversity conservation. Functional dissimilarities and similarities among species can be quantified by calculating pairwise functional distances between species, derived from differences in species’ traits (Violle et al. 2017). Functional dissimilarity promotes resource partitioning and coexistence, enhancing functional diversity and contributing to stronger, more stable ecosystem functioning over time. In contrast, functional similarity reflects the overlap in species’ functions, indicating potential ecological interchangeability (Eisenhauer et al. 2023). While often described as functional ‘redundancy’, such overlap can serve as an ecological insurance mechanism under environmental change, buffering the loss of species and ecosystem functioning (the insurance hypothesis; Yachi and Loreau 1999), and thus contributing to ecosystem resilience (Eisenhauer et al. 2023). Therefore, linking biodiversity to ecosystem functioning requires a multifaceted approach that goes beyond taxonomic diversity (Cadotte et al. 2009, Bae et al. 2018, Bełcik et al. 2020). Functional diversity, measuring differences in species traits, and phylogenetic diversity, measuring distances in evolutionary relatedness, both reflect ecosystem resilience and stability (Cadotte et al. 2011, 2012, Mori et al. 2013). Phylogenetically distant species can enhance ecosystem functioning through niche complementarity, which is often reflected in their traits (Cadotte 2013, 2017). Since many traits exhibit phylogenetic signals, meaning closely related species share similar traits (Cadotte et al. 2019), phylogenetic diversity can be used as a surrogate for unmeasured but conserved traits (Rothacher et al. 2025). Thus, functional and phylogenetic diversity often respond similarly to environmental change, but not necessarily similarly to taxonomic diversity (Bae et al. 2018).

Bats and birds are ideal model taxa to study biodiversity responses to habitat change. As mobile, higher trophic-level organisms, they respond sensitively to both abrupt and gradual environmental disturbances, including bottom-up processes, such as declines in prey availability (Russo et al. 2021). Their ecological role as predators is crucial for maintaining ecosystem stability (Terborgh et al. 2006), e.g., by controlling herbivorous insects and plant damage (Mooney et al. 2010, Böhm et al. 2011), and their protection under European law makes them highly conservation-relevant taxa (European Union 1992, 2009). Bats and birds exhibit diverse morphological, behavioral, and ecological traits that enable them to colonize various forest habitats and to navigate through the complex three-dimensional structure of forest environments (Jung et al. 2012, Froidevaux et al. 2016). With their diversity strongly linked to forest structure (Bradbury et al. 2005, Heidrich et al. 2020), they particularly benefit from heterogeneous forests, whether at early or late successional stages, which typically include canopy gaps and deadwood structures (Hilmers et al. 2018, Braunisch et al. 2019, Hendel et al. 2023, Jung et al. 2025). Forest bats and birds are associated with deadwood for roosting, nesting, and foraging (Bouvet et al. 2016, Kortmann et al. 2018, Wild et al. 2025), as well as with general stand characteristics, such as canopy openness, basal area, or vegetation height (Renner et al. 2018, Rigo et al. 2024). These responses are further modulated by foraging strategy and spatial scale, for example, open-space foraging bats cover larger areas than gleaning species (Tews et al. 2004, Hendel et al. 2023). Thus, forest structure can shape diversity patterns differently across spatial scales. Schall et al. (2018), for instance, found higher α-diversity of birds in fine-grained, uneven-aged forests in line with the early work by MacArthur and MacArthur (1961), whereas γ-diversity increased in coarse-grained forests of even-aged stands of different stages.

We experimentally enhanced forest structure in homogenized temperate forests by manipulating canopy cover and deadwood structures (Müller et al. 2023). Using a paired design across 11 sites and 234 forest patches, we compared structurally heterogeneous to homogeneous mixed beech (*Fagus sylvatica*) production forests, the dominant forest type in Germany. We assessed how enhanced structural heterogeneity affects multiple diversity facets of bats and birds, with emphasis on the responses of infrequent, frequent, and highly frequent species across different spatial scales.

Specifically, we tested three hypotheses:

1. γ-diversity of bats and birds is higher in structurally heterogeneous than in homogeneous forests, due to a greater variety of available niches.
2. Structural heterogeneity at the landscape level (γ-diversity) is mediated by varying contributions of α- and β-diversity, reflecting mechanisms of local species gain or dissimilarity in assemblages among sites.
3. Taxonomic diversity responds differently from phylogenetic and functional diversity in bats and birds, reflecting whether gained species are functionally and evolutionarily similar or dissimilar to the control assemblages.

We present a large-scale, spatially explicit (site-specific), and well-replicated field experiment to investigate the effects of forest heterogenization on biodiversity. Using autonomous data collection and automated species identification, a novel meta-analytic framework for paired experimental designs, and multifaceted diversity metrics, we show that γ-diversity increased in structurally enhanced forests for both bats and birds, but through distinct mechanisms. These results add new experimental evidence to the scientific debate on the effects of anthropogenic homogenization on diversity across scales, with key implications for forest restoration.

## Results

We identified a total of 17 bat species from 936 bat call recordings and 72 bird species from 16,147 bird sound recordings across all sites. As autonomous recorders do not provide true abundance data, we generally used an incidence-based approach for the analyses (Kortmann et al. 2025). To equalize sample completeness and avoid potential biases within both taxonomic groups, bat assemblages were standardized to the 25th quantile of sample coverage across all samples, extrapolated to double size: 0.819 (TD and PD) and 0.762 (FD), and for birds to 0.991, the minimum sample coverage across all samples when each is extrapolated to double size. Taxonomic (TD), phylogenetic (PD), and functional diversity (FD) are expressed in the same units of species, lineage, or functional group equivalents and can be directly compared within each diversity level (α, β, γ). Across levels, comparisons are only meaningful between α and γ, as multiplicative β-diversity is a ratio and represents how much of γ-diversity comes from differences between patches. Across all forest landscapes, bat γ-diversity consistently increased from homogeneous control to heterogeneous treatment forests across all diversity facets (TD, PD, FD) and diversity orders, emphasizing infrequent (q0), frequent (q1), and highly frequent (q2) species, lineages, and functional groups (Fig. 1, A). For birds, significant increases in γ-diversity were observed for TD and FD across all diversity facets and orders (Fig. 1, B). In bats, gains in γ-diversity were primarily driven by increased β-diversity (Fig. 1, C & E), whereas in birds, α-diversity was the only driver for γ (Fig. 1, D & F). Gains of TD for bats were strongest in γ- and β-diversity across all diversity orders, exceeding gains in PD and FD (Fig. 1, A & C). In contrast, birds showed the largest increases in γ- and α-diversity for FD at q1 and q2, exceeding the increases of TD (Fig.1, B & F).

**Figure 1.**
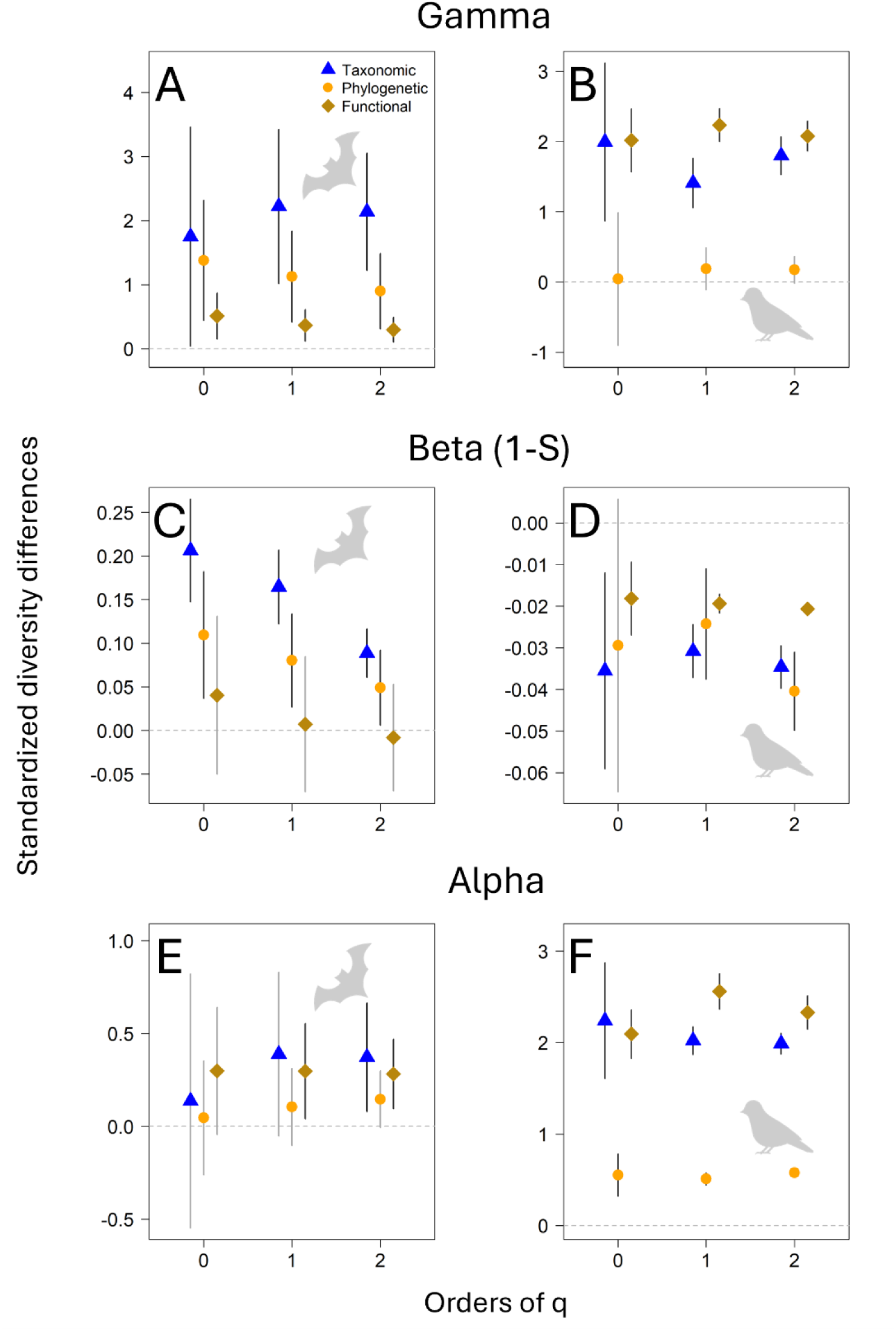
Results from three meta-analyses evaluating γ-diversity (A & B), β-diversity (C & D), and α-diversity (E & F) across 11 experimental forest landscapes. Symbols represent mean predicted changes in standardized diversity: blue triangles for taxonomic diversity (TD), orange circles for phylogenetic diversity (PD), and brown diamonds for functional diversity (FD). Error bars indicate 95% confidence intervals at fixed sample coverage levels. For bats, the level was set to the 25th quantile of sample coverage across all samples extrapolated to double size: 0.819 (TD and PD) and 0.762 (FD). For birds, it was set to 0.991, the minimum sample coverage across all samples when each is extrapolated to double size. The standardized diversity differences were calculated between each pair of control and treatment forest districts, across all three diversity facets (TD, PD, FD), and diversity orders referred to as q0, q1, and q2. Positive values indicate increased diversity in structurally heterogeneous forests relative to homogeneous controls. Differences are considered statistically significant when the confidence interval does not include zero, as indicated by dark-colored error bars. TD, PD, and FD are expressed in the same units of species, lineage, or functional group equivalents and can be directly compared within each diversity level (α, β, γ). To account for variation in patch numbers between sites, we applied a Jaccard-type turnover transformation (1-S) of multiplicative β-diversity, which quantifies dissimilarity between assemblages relative to γ (see methods). Across levels, only α and γ can be directly and meaningfully compared (e.g., for q0: TD α < TD γ). For detailed results, see forest plots in the supplements (Document S1. Fig. S3-29 and Fig. S30-56).

### γ-diversity increased across heterogeneous forest landscapes for bats and birds

For bats, taxonomic γ-diversity increased significantly in structurally enhanced landscapes with gains of approximately two effective numbers of species across all diversity orders (q0 = 1.8, q1 = 2.2, q2 = 2.1) (Fig. 1, A). These were the strongest gains among the three diversity facets, exceeding those observed for PD (q0 = 1.4, q1 = 1.1, q2 = 0.90) and FD (q0 = 0.51, q1 = 0.37, q2 = 0.30).

Birds also showed taxonomic gains of about two effective species (q0 = 2.0, q1 = 1.4, q2 = 1.8) (Fig. 1, B). FD showed the strongest gains in γ-diversity for increasingly frequent groups (q0 = 2.0, q1 = 2.2, q2 = 2.1).

### β-diversity increased for bats but decreased for birds across treatment forest districts

Bats showed increases in taxonomic (q0 = 0.21, q1 = 0.17, q2 = 0.089) and phylogenetic (q0 = 0.11, q1 = 0.080, q2 = 0.049) β-diversity, for all orders of q, with effect sizes declining from infrequent to highly frequent species (Fig. 1, C).

In contrast, birds showed overall significant decreases in β-diversity for taxonomic (q0 = −0.036, q1 = −0.031, q2 = −0.035) (Fig. 1, D), phylogenetic, except at q0 (q1 = −0.024, q2 = - 0.040), and functional diversity (q0 = −0.018, q1 = −0.019, q2 = −0.021).

### α-diversity showed stronger gains for birds than for bats in treatment patches

Bats showed significant increases in taxonomic α-diversity among highly frequent species (q2 = 0.37) (Fig. 1, E). Functional diversity increased significantly for frequent and highly frequent groups (q1 = 0.30, q2 = 0.28).

Birds showed consistent significant gains in taxonomic α-diversity, with gains of about two effective species (q0 = 2.2, q1 = 2.0, q2 = 2.0) (Fig. 1, F). Phylogenetic α-diversity increased significantly across all orders (q0 = 0.55, q1 = 0.51, q2 = 0.58), but with smaller effect sizes than for FD, which showed the strongest gains for increasingly frequent functional groups (q0 = 2.1, q1 = 2.6, q2 = 2.3).

## Discussion

### Increased γ-diversity in heterogeneous forest landscapes is driven by distinct α- and β-diversity mechanisms for bats and birds

Overall, γ-diversity of bats and birds increased in structurally heterogeneous forests, except for birds’ PD, supporting our first hypothesis and reinforcing the habitat-heterogeneity hypothesis: small-scale interventions that enhance habitat complexity can promote biodiversity by expanding ecological niche availability. However, consistent with our second hypothesis, the underlying mechanisms driving γ-diversity differed, reflecting distinct contributions of α- and β-diversity for bats and birds.

In bats, increases in landscape-scale diversity are primarily driven by increased β-diversity, indicating greater dissimilarity of assemblages between patches with different light conditions and deadwood amounts. Local α-diversity gains of bats were small and only became apparent at the landscape scale, particularly for species with high occurrence frequency (q1, q2). This is consistent with the dominance of β-diversity hypothesis, which states that γ-diversity is driven by β-rather than α-diversity (Tscharntke et al. 2012). This pattern has been supported by other studies (Farnsworth et al. 2014, Morante-Filho et al. 2016, Costa and Schmidt 2022) and aligns with bats’ high mobility and wide-ranging foraging behavior. Their habitat use is strongly shaped by forest structure, with different foraging guilds specializing in distinct forest structures. This niche differentiation allows them to locate and exploit spatially distributed resources while reducing interspecific competition (Schnitzler and Kalko 2001, Müller et al. 2012, Charbonnier et al. 2014). Closed-space foragers (e.g., *Myotis bechsteinii, M. nattereri, Plecotus auritus*) prefer dense forest interiors or canopies. In contrast, edge-space foragers (e.g., *Barbastella barbastellus, Myotis myotis, Pipistrellus pipistrellus*) prefer more open forest stands with sparse understory and leaf-littered ground, and can exploit forest edges and gap margins. Open-space foragers (e.g., *Eptesicus* and *Nyctalus* spp.) hunt in open canopies or above gaps. In this study, homogeneous control forests with closed canopies mainly supported closed- and edge-space foragers. Aggregated treatments created canopy gaps and edge structures, likely attracting open-space foragers (Müller et al. 2013, Kortmann et al. 2018, Jung et al. 2025) (also see Document S1, Fig. S1, and Tab. S1). A study comparing forest patches with high and low structural complexity showed that TD, PD, and FD of aerial-hawking bats increased at habitat edges, highlighting the importance of heterogeneous landscape elements (López-Baucells et al. 2022). Distributed treatments with deadwood structures under closed-canopy likely favored closed-space specialists, adapted to navigate in dense habitats, which harbor high prey abundance (Müller et al. 2012, Hochrein et al. 2025, Jung et al. 2025). Additionally, all forest bats depend on structures associated with old or damaged trees for roosting, with species-specific preferences for cavities or crevices (Kortmann et al. 2018). Our treatments replicated these features across forest patches by varying deadwood volume and type, such as snags and habitat trees. Even lying deadwood volume and understory vegetation can enhance bat foraging activity, suggesting the influence of bottom-up processes (Jung et al. 2025). Social information transfer about new roosts (Kerth and Reckardt 2003) or food resources may have facilitated resource exploitation in heterogeneous forest landscapes (Safi and Kerth 2007), promoting differentiation in local bat assemblages across patches (increased β-diversity), especially among rarer, specialized species (β: TD & PD at q0).

In contrast, bird γ-diversity increased exclusively through temporal gains in within-patch α-diversity, despite declining β-diversity, underlining the effect of small-scale and often randomly distributed structural features. Although it has been suggested that opposing α- and β-trends might cancel each other out (Farnsworth et al. 2014), our results suggest that structurally heterogeneous forests can simultaneously enhance local and regional bird diversity while reducing assemblage differentiation among sites. Unlike bats, most forest bird species operate during the breeding season on smaller spatial scales within fixed territories that are actively and aggressively defended (Krebs 1982, Poesel and Dabelsteen 2005), limiting their spatial mobility and thus also potential changes in assemblages across patches. Deadwood and canopy gaps can locally and temporally increase insect availability (Lettenmaier et al. 2022, Rothacher et al. 2023, 2025), leading to a localized diversification of resources, as already shown at the scale of a few trees in the seminal work by MacArthur and MacArthur (1961). This increase in resources may have attracted infrequent species, such as rare cavity- or deadwood-associated specialists (α: TD at q0), and increased local species richness (Whittaker 1972). Yet, as the same rare species may respond similarly across sites, this can lead to increases in α- and γ-diversity without a corresponding rise in β-diversity. Similarly, increases also occurred among more common species (α: TD at q1 and q2), likely leading to the same species quickly occupying niche spaces across all patches, making assemblages more similar, and further reducing β-diversity. However, unlike our results, a study in heterogeneous mixed forests in Germany found no significant effects of structural features on bird α-diversity, but did find effects on β-diversity, particularly related to light availability, which likely increased insect resources and foraging space (Schauer et al. 2023). At early successional stages, forest gaps increase sunlight exposure, supporting herbaceous understory plants that produce seeds and fruits, providing resources for both omnivorous and herbivorous bird species. β-diversity effects may have decreased over time (Freitag Kramer et al. 2025), as positive effects of structural heterogeneity on diversity are typically strongest during the early and late successional stages (Hilmers et al. 2018). In our study, sampling occurred 4 to 7 years post-intervention, when forest succession had already reached an intermediate stage, with some patches showing intensive regeneration (University Forest).

### Diversity facet responses: bats show patterns of functional similarity, birds gain dissimilar traits

Bats showed greater increases in TD than in PD or FD at both β- and γ-scales (TD > PD, FD), supporting our third hypothesis, and suggesting that functionally similar species contributed the most to the detected diversity gains (Flynn et al. 2009). An increase of two effective species at standardized sample coverage is substantial for bats, given that only 25 species occur in Germany (Bundesamt für Naturschutz 2025). Unlike birds, bats are not highly territorial over feeding areas and can coexist despite exploiting similar resources by differing in foraging strategies and operating at large spatial scales. However, because bats can be classified into only a few broad guilds, this leads to trait overlap within guilds (Denzinger and Schnitzler 2013). Moreover, all observed bat species are closely related, belonging to the same family of Vespertilionidae, which is divided into a few subfamilies that broadly correlate with their foraging guilds, e.g., Myotinae (*Myotis*) occupy diverse niches, such as edge and closed spaces, and Vespertilioninae (e.g., *Nyctalus*, *Eptesicus*, *Vespertilio*, *Pipistrellus*) dominate open spaces.

Phylogenetic diversity patterns in birds, contrary to our third hypothesis, did not align closely with those of FD. Both diversity facets can capture complementary aspects about community patterns (Cadotte et al. 2013), e.g., functional and phylogenetic bird diversity can respond similarly to vegetation or landscape structure, but differently from taxonomic diversity (Klingbeil and Willig 2016, Bae et al. 2018). However, in our study, FD and TD responded more strongly to heterogeneous forest structure than PD (FD, TD > PD), suggesting that the gained species at the α-scale contributed only a few new phylogenetic lineages, but these did not translate into an overall increase in PD at the γ-scale. Over 60% of the observed bird species belonged to the species-rich order of Passeriformes, most of which were widely distributed and occurred in both control and enhanced forests (see Document S1, Fig. S2). Only a few species were unique to the enhanced forests, adding a new lineage: the rare ring ouzel (*Turdus torquatus*), and the green sandpiper (*Tringa ochropus*), a small wader, breeding in wet forests, introducing an entirely new order to our phylogenetic tree. Although PD and FD are often correlated, they are not interchangeable. Phylogenies and traits may capture different ecological dimensions: phylogeny reflects deeper, ancestral similarities, while traits may highlight more recent evolutionary changes (Cadotte et al. 2019). For example, this is highly pronounced in the functional dissimilarity of bat species in the genus *Myotis* (Document S1, Fig. S1, Tab. S1). The closer alignment of FD with TD, relative to PD, points to short-term ecological processes driving species gains at the patch level (Aguirre et al. 2016), e.g., temporal increase in food resources or nesting sites to which birds quickly respond. Among increasingly frequent species (at q1 and q2) of α- and γ-diversity, gains in FD were higher than in TD (FD > TD), suggesting that the added species contributed dissimilar or complementary ecological functions (Moreno et al. 2024). As birds are territorial, increases in effective species richness were modest at the local scale (∼2 species). However, these gains added 2–2.5 new functional groups among more frequent species, suggesting greater functional diversification without increasing competition (Moreno et al. 2024).

### Diversity order patterns: infrequent species exhibit variable responses, increasingly frequent species respond uniformly

Within each diversity level (α-, β-, and γ), response patterns remained relatively consistent across diversity orders (q0–q2) for both bats and birds. However, diversity differences between control and treatment forests were more variable at q0, while effect sizes were often stronger and more stable at q1 and q2, suggesting a more heterogeneous response among infrequent species (q0), while increasingly frequent species (q1, q2) responded more uniformly to structural changes. These trends mirror findings from a bird sound study in Ecuador, which also found more variance in diversity values for q0 (Kortmann et al. 2025). However, dissimilarity-based methods have been criticized for producing misleading results, when species are undersampled and dissimilarity values become saturated when many sites share no species, which is often the case for sampling rare species, emphasized at q0 (Tuomisto et al. 2012). Our standardization of sample coverage within each taxa mitigates this sampling bias to some extent. Furthermore, as we used acoustic data, which provides incidence-based data, we lacked information on species abundances reflecting their dominance within assemblages (Tucker et al. 2016). Yet, this was also addressed by our method, by using repeated measurements at each patch and analyzing our data on a nightly basis for bats and a daily basis for birds, we increased the number of sampling units.

### Conservation and restoration implications

Environmental heterogeneity, particularly structural complexity, is strongly associated with increased species diversity across spatial scales (Stein et al. 2014). Our study supports this link, showing that restoring structural heterogeneity in production forests can promote bat and bird diversity, however, effects vary across spatial scales and taxa (Schall et al. 2018, Heidrich et al. 2020, Schauer et al. 2023).

For bats, enhancing the spatial variation of structural heterogeneity at broader scales is essential. Treatments that create contrasting canopy conditions (open vs. closed) increase horizontal structural diversity and microclimate variation, which was confirmed on the same patches by other studies (Thom et al. 2020, Pierick et al. 2025), altering understory vegetation and prey availability (Hendel et al. 2025). For example, moths are often associated with closed canopies, while beetles and other arthropods, both saproxylic and non-saproxylic, depend on deadwood (Seibold et al. 2016, Hendel et al. 2025). Diverse deadwood structures, e.g., lying, standing, and different decay stages, provide diverse microhabitats for both arthropods and roosting bats (Tillon et al. 2016, Seibold et al. 2023, Rothacher et al. 2023), promoting assemblage differentiation between stands and increasing bat diversity at the landscape scale (Barnes et al. 2016, Jung et al. 2025).

In contrast, birds responded positively to increased heterogeneity regardless of spatial distribution. Opening closed forest canopies can promote forest specialists and generalists, enhancing overall γ-diversity (Vanderwel et al. 2007, Schall et al. 2018). In forests near cultural landscapes, forest gaps can attract some opportunistic birds of open habitats (Żmihorski et al. 2016) such as the Yellowhammer (*Emberiza citrinella*), the Common linnet (*Linaria cannabina*), or the European turtle dove (*Streptopelia turtur*), as observed on our experimental patches located in some regions (e.g., Bavarian Forest, University Forest, and Lübeck). Furthermore, creating gaps initiates forest succession, enhancing vertical structural diversity and supporting overall bird diversity (Lesak et al. 2011, Vogeler et al. 2014, Hanle et al. 2020), particularly shrub-nesting birds and short-distance migrants (Graser et al. 2025). Additionally, deadwood structures can serve as nesting sites, foraging substrates, and singing perches (Bouvet et al. 2016). For example, snags within small forest gaps can expand territories and promote bird diversity in deciduous forests, especially for cavity-nesting species that arrive relatively late from their wintering grounds (Müller 2005, Lewandowski et al. 2021).

Effective management practices should promote structural heterogeneity at the local and landscape scales. This can be achieved through a benign neglect strategy in large, protected areas, which deliberately allows natural disturbances to occur, thereby creating heterogeneous landscapes with suitable structures for bats and birds (Müller et al. 2010, Kortmann et al. 2018, Cours and Duflot 2025). Based on our results, we suggest to locally promote gaps and deadwood structures whenever possible to enhance bird diversity in production forests. Moreover, combining different silvicultural approaches, e.g., gap felling, shelterwood, or single-tree selection, applied with varying intensity in a spatially explicit design (Schall et al. 2018) will foster habitat heterogeneity and benefit both birds and bats. In turn, supporting bats and birds enhances natural pest control, contributing to healthy and resilient forest ecosystems (Mäntylä et al. 2011, Blažek et al. 2021).

### Study limitations

Some site-specific deviations from general patterns were observed, but our meta-analytic approach minimized their influence through appropriate weighting. For example, the unrealistic high estimate for taxonomic α-diversity of bats at a single study site in Hunsrück (TD q0 in control = 35.8) had minimal influence on the overall outcomes (weight = 0.05%). Furthermore, diversity estimates were based on incidence data from acoustic recordings, reflecting presence or activity instead of absolute abundances, while this limits direct interpretation of species counts, it provides robust, standardized insight into acoustic community structure (Kortmann et al. 2025). However, this approach also requires a sufficient number of sampling units per patch to ensure analytical robustness. Thus, we recommend using at least eight sampling units per patch. To test the robustness of observed patterns, meta-analyses can be repeated with varying sample coverage levels. Consistent results strengthen the confidence in inferences, e.g., using the median of sample coverage across all samples extrapolated to double size, yielded the same patterns (see bat analysis with SC threshold = C50% in Supplementary Document S1, Fig. S57). Our study focused on assessing spatial differences in species assemblages, based on a one-year sampling campaign at an intermediate stage of forest succession. This approach allowed species enough time to redistribute among patches after the interventions, however, we could not analyze temporal changes in assemblage, which can fluctuate over time (Rolls et al. 2023). This should be considered in future studies.

## Conclusion and outlook

Global biodiversity is undergoing significant changes, with α- and β-diversity playing distinct roles in shaping γ-diversity (Gonçalves-Souza et al. 2025, Keck et al. 2025). In this study, we experimentally enhanced structural between-patch complexity in homogenized temperate production forests by modifying canopy cover and deadwood availability. Our results confirmed that (1) γ-diversity of bats and birds is higher in structurally heterogeneous than homogeneous forest landscapes, except for birds’ PD, and (2) γ-diversity is driven by distinct contributions of α- and β-diversity in each group. Bats primarily respond to spatial variation across the landscape (β-diversity), consistent with their high mobility and foraging range, while birds respond mainly through local species gains (α-diversity), influenced by opportunistic behavior and competition. In line with our third hypothesis, (3) TD increased more strongly than PD and FD in heterogeneous forests for β- and γ-diversity of bats, suggesting functional similarity in gained species. In contrast, for birds, FD aligned more closely with TD than PD, indicating that newly gained bird species were functionally dissimilar to the control assemblages. These contrasting responses underscore the importance of taxon-specific approaches in biodiversity-oriented forest management, even within the same trophic level. By contributing to a broader understanding of how α- and β-diversity respond to habitat heterogeneity, our study gives critical insights for developing effective, scale-sensitive restoration strategies.

## Resource availability

## Lead contact

Requests for further information and resources should be directed to and will be fulfilled by the lead contact, Clara Wild.

## Materials availability

This study did not generate new unique reagents.

## Data and code availability

● Bat species, trait, and phylogenetic data have been deposited at Zenodo and are publicly available as of the date of publication at [DOI].
● Bird species, trait, and phylogenetic data have been deposited at Zenodo and are publicly available as of the date of publication at [DOI].
● All original code has been deposited at Zenodo and is publicly available at [DOI] as of the date of publication.
● Any additional information required to reanalyze the data reported in this paper is available from the lead contact upon request.

## Acknowledgments

We thank all local managers for their support in our research. In particular, we thank K. Kraus, A. Kieffer, G. Bach, J. Torres, K. Kallnik, M. Mauermann, T. Becker, T. Rosch, W. Weisser, the interns of the Bavarian Forest National Park, and all other assistants. We also thank J. Thein and C. Franz for providing bat recorders, and R. Martin for validating our bird sound data. The experimental sites were established within the project ‘Beta-Diversität experimentell für nachhaltige Waldbewirtschaftung in Mitteleuropa’ of the German Federal Environmental Foundation (DBU) (project no. 34488/01) and the BioHolz project (grant no. 01LC1323A). CW, OM, JR, ST, SK, and JM received funding from the German Research Foundation (DFG) within the research unit ‘BETA-FOR’ (project no. 459717468). JR was further supported by funding from the Bavarian State Ministry for Food, Agriculture, and Forestry (StMELF) (grant no. L062). MK acknowledges funding by the Bavarian Research Institute for Digital Transformation (bidt), an institute of the Bavarian Academy of Sciences and Humanities (ROOT: Real-time earth Observation of fOrest dynamics and biodiversiTy (KON-22-024)).

## Author contributions

Conceptualization: JM, CW; Data curation: CW, SK, OM; Formal analysis: CW, JM, OM; Funding acquisition: JM, ST; Investigation: CW, JR, RP, RH, KS, SH, SM, MJ, ND, JS; Methodology: AC, OM, SK, MK; Project administration: CW, JR, RP, OD; Resources: JM; Software: AC, PYC, SK, OM; Supervision: JM, ST, MC; Validation: JM, CW, OM; Visualization: CW, MK; Writing – original draft: all authors; Writing – review & editing: all authors.

## Declaration of generative AI and AI-assisted technologies

During the preparation of this work, the authors used ChatGPT and Grammarly in order to improve the orthography, grammar, and writing style. Additionally, the authors used the AI-based tools BirdNET and batIdent to assist in the identification of bird and bat species. After using these tools, the authors reviewed and edited the content as needed and take full responsibility for the content of the published article.

## Supplemental information

Document S1. Figures S1–S57 and Table S1–S4

Table S2. Excel file containing additional data too large to fit in a PDF, related to Figure 2

## Methods

### Study area and experimental design

**Figure 2.**
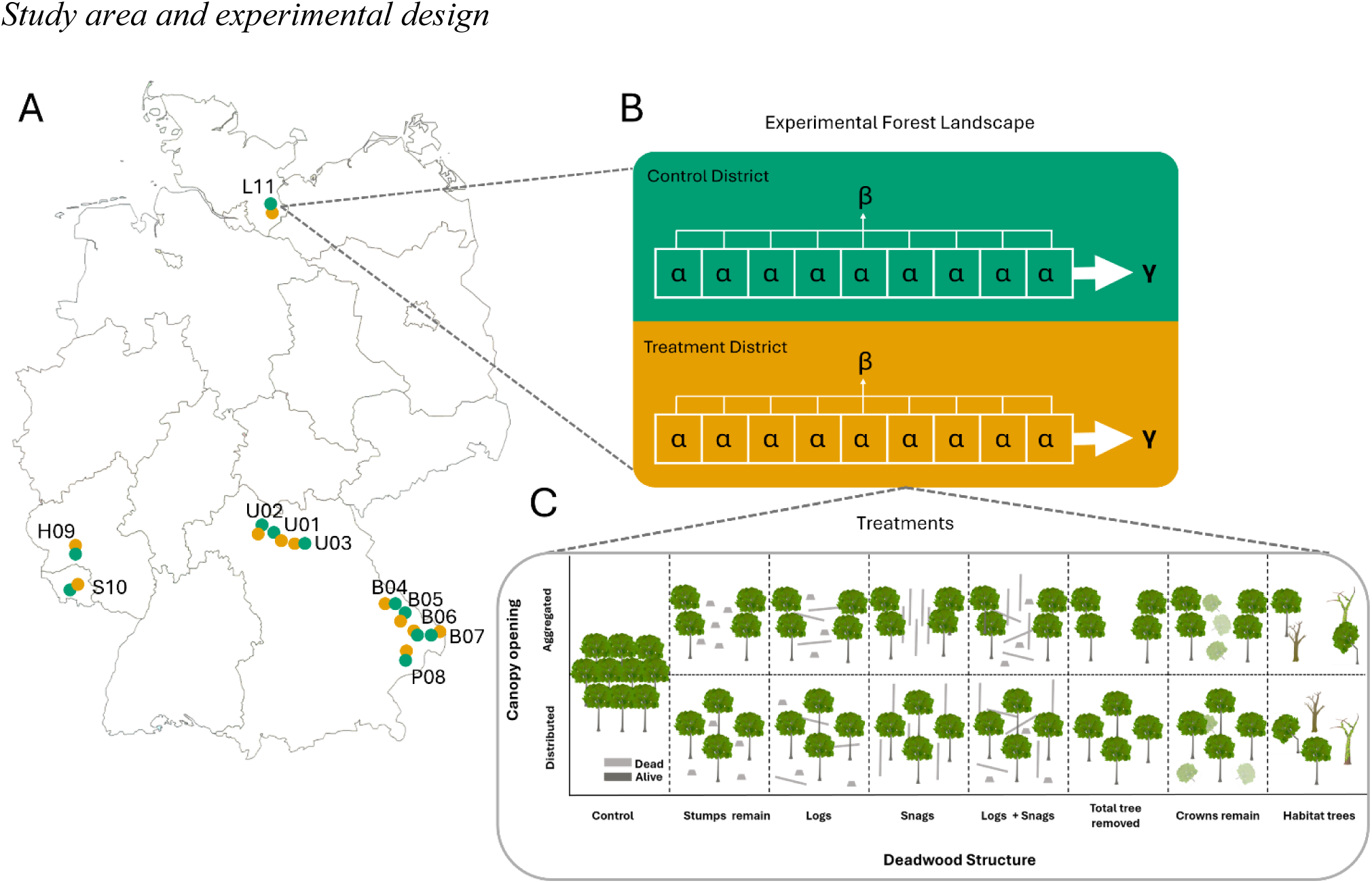
Experimental design of the BETA-FOR research unit. (A) Map of Germany showing six study regions with 11 experimental forest landscapes: Lübeck (L11), Saarland (S10), University Forest (U01–U03), Passau (P08), Hunsrück-Hochwald National Park (H09), and Bavarian Forest National Park (B04–B07). Each landscape includes two paired forest districts: a treatment district with enhanced structural heterogeneity (orange), and a control district representing homogeneous production forest (green). (B) Schematic illustration of an experimental landscape, illustrating spatial scales and diversity levels assessed: each district contains 9 patches (15 in the University Forest), used to evaluate within-patch α-diversity, between-patch β-diversity, and overall γ-diversity. (C) Overview of 14 treatment types and a control patch. The depicted control patch serves as local control within each treatment district and further represents the stand structure of all patches in the control district. Treatment patches, established only in treatment districts, vary in spatial arrangement, either aggregated, within a forest gap, or distributed, under a closed canopy. The control patch and the first eight treatments from left to right (Stumps remain, Logs, Snags, Logs + Snags) are replicated across all experimental landscapes, while six additional treatments (Total tree removed, Crowns remain, Habitat trees) are exclusive to the University Forest. Adapted from a graph provided by Rike Schwarz.

This study is part of the collaborative research unit BETA-FOR (Müller et al. 2023). The study design comprises 234 experimental patches distributed across 11 forest landscapes in six regions of Germany, spanning a gradient of mean elevation above sea level (a.s.l.): Lübeck (41 m), Saarland (296 m), the University Forest of the Julius-Maximilians-Universität Würzburg (336 m), Passau (496 m), the Hunsrück-Hochwald National Park (654 m), and the Bavarian Forest National Park (931 m) (Fig. 2, A; for detailed site information, see supporting information, Table S2. Excel file). All forests are managed and reflect the typical tree composition of Central European mixed forests. These are primarily dominated by deciduous tree species, with conifers comprising 1% to 30% of the total basal area per forest. European beech (*Fagus sylvatica*) is the dominant species in five of the regions, while the University Forest features a mixed composition of beech, maple (*Acer spp.*), ash (*Fraxinus excelsior*), hornbeam (*Carpinus betulus*), and oak (*Quercus spp.*).

Each experimental landscape consists of two paired forest districts: the treatment district is characterized by enhanced structural heterogeneity (heterogeneous forest), and the control district represents an untreated production forest (homogeneous forest). This design allows for contrasting findings to a reference control and to study biodiversity at different scales (Keck et al. 2025). Within each forest district, nine patches, and 15 in the University Forest, were established, each measuring 50 m × 50 m. This design enables assessment of within-patch α-diversity, between-patch β-diversity, and overall γ-diversity within each district (Fig.2, B).

Structural enhancements in heterogeneous forests increased between-patch complexity by assigning each patch within a treatment district a distinct combination of canopy openings and deadwood structures. These treatments were not repeated within districts but were replicated across the 11 experimental landscapes, except for six treatments implemented only in the University Forest. Aggregated treatments created canopy gaps with diameters of 25–30 m, whereas distributed treatments maintained a closed canopy. These interventions resulted in microclimatic contrasts, with forest gaps exhibiting higher UV radiation and greater temperature fluctuations, warmer during the day and cooler at night, compared to the more thermally buffered conditions under closed canopies (Thom et al. 2020). Deadwood structures, such as stumps, snags, logs, and combinations thereof (with additional elements in the University Forest), were established both in canopy gaps and under closed canopies (Fig. 2, C). Patches and treatments were established in the winters of 2015/16 (Bavarian Forest and Passau), 2016/17 (Saarland, Lübeck, and Hunsrück), and 2018/19 (University Forest), allowing sufficient time for colonization by forest fauna.

### Bat sampling and data preparation

Due to the wide geographic distribution of study sites across Germany, sampling was conducted in 2022 in the University Forest, Bavarian Forest National Park, and Passau regions and in 2023 in the Saarland, Hunsrück, and Lübeck regions. Bats were surveyed using autonomous bat recorders (batcorder, generations 2 and 3; ecoObs GmbH, Nürnberg, Germany), which recorded echolocation calls for at least one night per patch and month in May, June, and July. Additional sampling nights in May, June, and August were included to compensate for adverse weather or other suboptimal sampling conditions. The limited availability of recording devices restricted sampling to one pair of forest districts per night. One batcorder was installed in each patch within a 15-meter radius of the patch center, mounted at a height of 1.50 to 1.70 m on the trunk of a thin tree, leafless at recorder height to minimize background noise. The antenna was angled upward toward the patch center or, when available, toward structural features such as canopy gaps or skid trails, which are often used by bats for navigation. Batcorders were configured with the following settings: recording quality set to 20, post-trigger duration of 400 ms, threshold level at −27 dB, and a critical frequency of 16 kHz. Recording was continuous and set to start one hour before sunset and end one hour after sunrise. Acoustic data were processed using bcAdmin4 (Version 1.2.6, ecoObs GmbH) to organize and analyze call sequences, followed by species-level identification using batIdent (Version 1.5, ecoObs GmbH), following the approach in Müller et al. (2013). The minimum detection probability was set to 0.6. For subsequent analyses, only species-level identifications were considered, with the exception of *Myotis brandtii/mystacinus* and *Plecotus spp.*, which cannot be reliably distinguished acoustically. However, for the assignment of trait data and phylogenetic information, these groups were attributed to *Myotis mystacinus* and *Plecotus auritus*, respectively. Additionally, species not known to occur in the respective study regions were excluded based on expert knowledge and verification of individual audio files. Raw data were initially expressed as ‘minute counts’, defined as the number of minutes within a recording session during which at least one bat call was detected (Hochrein et al. 2025). Since individuals cannot be reliably identified from acoustic recordings, we converted these data to incidence-based presence– absence format (0 = no detection, 1 = detection) for each sampling night (‘sampling unit’ for bats; Document S1, Tab. S3).

To construct a phylogenetic tree, species-specific information was extracted by subsetting the mammalian megatree published by Upham et al. (2019). A customized phylogeny was generated using the ‘phylo.maker’ function from the *U.PhyloMaker* package, following the workflow provided in Jin and Qian (2023). The resulting tree was saved in *.nwk* format. For visualization, we used the ‘ggtree’ function from the *ggtree* package (Yu 2022), following the approach described by Kortmann et al. (2025) (Document S1. Fig. S1).

We selected seven traits that represent morphological, behavioral, and functional adaptations of bats related to using forest structures (Document S1., Tab. S1). Trait data for each species were primarily obtained from the EuroBaTrait 1.0 dataset (Froidevaux et al. 2023) and supplemented with information from Müller et al. (2012, 2013). All morphological traits were log_e_-transformed. For Ear length and Tail length, we calculated residuals from a linear model with the ‘lm’ function (R Core Team 2024) using log_e_-transformed Forearm length as the predictor to account for body size scaling. In addition, log_e_(Body mass), Call duration, and Maximum frequency were standardized (mean = 0, SD = 1) using the ‘decostand’ function (method = standardize) from the *vegan* package (Oksanen et al. 2022).

### Bird sampling and data preparation

Bird sampling, similar to bat surveys, was conducted in 2022 in the University Forest, Bavarian Forest National Park, and Passau regions, and in 2023 in the Saarland, Hunsrück, and Lübeck regions. Birds were sampled using autonomous sound recorders (BAR and BAR-LT; Frontier Labs, Australia), which recorded ambient sound daily from March to June. One recorder was installed within a 15-meter radius of the patch center, mounted on the trunk of a thin tree, leafless at recorder height to optimize recording quality. Recorders were placed at a height of 1.50 to 1.70 m with the microphone facing downward. Recorders were set to record for 2 minutes every 12 minutes, using a sampling rate of 44.1 kHz. Recordings were scheduled daily to begin two hours before sunrise and continue until four hours after sunrise, and again from three hours before sunset to three hours after sunset. For subsequent analyses, the recording period was restricted to March 24 to June 15, coinciding with the peak breeding and singing activity of most bird species. Sound data were analyzed using BirdNET version 2.4. We extracted all identifications with a confidence value above 0.5. In a second step, ∼15,000 expert-validated species identifications from our study sites were used to calculate a species-specific threshold, following the method of Wood and Kahl (2024). For some rare species, we applied a strict confidence threshold of 0.95 (Seibold et al. 2024). To achieve the same level as for bats (one-night campaigns), we aggregated the bird data on a daily basis, with each species counted only once per daily recording period. The resulting dataset was compiled in an incidence-based presence–absence format (0 = no detection, 1 = detection) for each sampling day (‘sampling unit’ for birds; Document S1, Tab. S4). The final data set was once more checked by JM for plausibility of species occurrences in the study regions, and implausible species were verified and, if erroneous, removed.

Phylogenetic information was obtained from the bird megatree (Jetz et al. 2012) and adjusted to the observed species using the same procedure applied to bats, employing the *U.PhyloMaker* package (Jin and Qian 2023). The resulting tree was saved in *.nwk* format and visualized using the ‘ggtree’ function from the *ggtree* package (Yu 2022) (Document S1., Fig. S2).

Similar to the approach used for bats, we selected seven bird traits from the AVONET dataset (Tobias et al. 2022) that reflect adaptations to forest structure use (Document S1., Tab. S2). All morphological traits were log_e_-transformed. For Beak length, Tarsus length, and Tail length, we calculated residuals from linear models using the ‘lm’ function (R Core Team 2024), controlling for log_e_-transformed Body mass. Additionally, log_e_-transformed Body mass was standardized (mean = 0, SD = 1) using the ‘decostand’ function (method = “standardize”) from the *vegan* package (Oksanen et al. 2022).

### Data analysis

All analyses were conducted using R version 4.4.2. To assess diversity differences between treatment (structurally heterogeneous) and control (homogeneous) forest districts, we applied a comparative meta-analytic approach. This approach combines two core concepts, embedded into the iNEXT.3D framework (Chao et al. 2021): sample coverage, to standardize sample completeness, and Hill numbers, to unify diversity measurements across different orders of q, including species richness (q = 0), Shannon diversity (q = 1), and the Simpson diversity (q =2). These tools provide robust, comparable biodiversity estimates across sites (Roswell et al. 2021, Kortmann et al. 2025).

As our data is based on incidences (presence-absence), and not abundances, we use Hill-Chao numbers (q = 0, 1, 2; hereafter ‘diversity orders’), which extend the original Hill number approach, developed for taxonomic diversity (TD), to phylogenetic (PD) and functional diversity (FD) (hereafter ‘diversity facets’) and incidence-based occurrence data (Chao et al. 2021). For bats, only four sampling units (nights) per patch were available, which can be insufficient for reliable inference using incidence-based models, particularly for FD (Colwell and Chao 2022) (see Document S1, Tab. S3). To address this for FD, we used species incidences summed per patch as a “detection frequency”, serving as a proxy for abundance frequency. The outputs for FD of bats were generated using abundance-based models, including the estimation of sample coverage. TD quantifies the effective number of species and represents species richness, whereas PD quantifies the effective number of lineages, and FD quantifies the effective number of virtual functional groups, or functional species, both expressing ecosystem functioning and resilience (Cadotte et al. 2012, Chao et al. 2021). Diversity orders capture the relative frequency of species incidence within the assemblage, emphasizing infrequent (q0), frequent (q1), and highly frequent (q2) species (Kortmann et al. 2025).

To calculate the functional diversity, a species-pairwise distance matrix is needed. Thus, we computed Gower’s distance (Gower 1971) between species for bats and birds based on their traits, using the function ‘daisy’ from the *cluster* package (Maechler et al. 2024). Patches with no observed species were excluded from the analysis, reducing the number of patches in some districts. Due to malfunction of recorders or adverse weather conditions, recording had to be repeated occasionally. This resulted in varying numbers of sampling units per patch, which are defined as one recording night for bats and one recording day for birds (Document S1, Tab. S3 and S4). In some cases, the number of patches differed between treatment and paired control districts (e.g., E = 8 vs. C = 9 patches), which can influence the range of β-diversity values. Thus, a Jaccard-type turnover (1-S) transformation was applied to β-diversity (for details, see Tab. 1 in Chao et al. 2019). This metric quantifies dissimilarity between assemblages relative to γ-diversity, standardized by patch number, with a value of zero indicating identical assemblages in the patches and one indicating no shared species. To account for sample incompleteness within the meta-analytic framework, sample coverage (SC) was estimated using the function ‘Coverage’ from the iNEXT.3D package (Chao et al. 2021), setting the minimum quantile across all SC(2n) values to be greater than 0.5. At the fixed sampling coverage level, rarefaction and extrapolation curves were generated for each of the eleven treatment-control district pairs.

Using the ‘iNEXTmeta_beta’ function (Chao 2025), landscape-level γ-diversity was decomposed into its components of within-patch α-diversity and between-patch β-diversity. The three diversity facets (TD, PD, and FD) were calculated for each district pair and diversity level across all three diversity orders, referred to as q0, q1, and q2 (Chao et al. 2023a). All metrics are expressed in the same units of species, lineage, or functional group equivalents and can be directly compared within each diversity level (α, β, γ) (Chao et al. 2021). Across levels, comparisons are only meaningful between α and γ, as multiplicative β-diversity is a ratio, representing the effective number of non-overlapping assemblages, calculated based on a multiplicative diversity composition (β = γ/α). Thus, α-diversity captures temporal changes of the within-patch assemblage, β-diversity reflects spatial differentiation among assemblages, and γ-diversity integrates the spatio-temporal variation of communities across all study sites.

For each treatment-control district pair, we calculated diversity differences, with positive values indicating higher diversity in heterogeneous forests compared to homogeneous forests. The overall effect across all 11 experimental landscapes was estimated through a meta-analysis (see Data and code availability). Confidence intervals (CI) and average differences between all forest pairs were calculated using bootstrapping (nboot = 50). Significance is inferred when the CI does not include zero. For reproducibility, all models were run with ‘set.seed(1)’ or ‘clusterSetRNGStream(cl, iseed = 1)’ for parallel computation of PD with ‘parLapply’ from the package *parallel* (R Core Team 2024). Forest plots illustrating the standardized diversity differences were generated using ‘ggiNEXTmeta’(Chao 2025) and are presented in the supplements (Document S1., Fig. S3–29 and Fig. S30-56) (Chao et al. 2023a, b).

