## Supplemental information for "Higher bat and bird γ-diversity in structurally complex forests is driven by distinct α- and β-diversity responses"

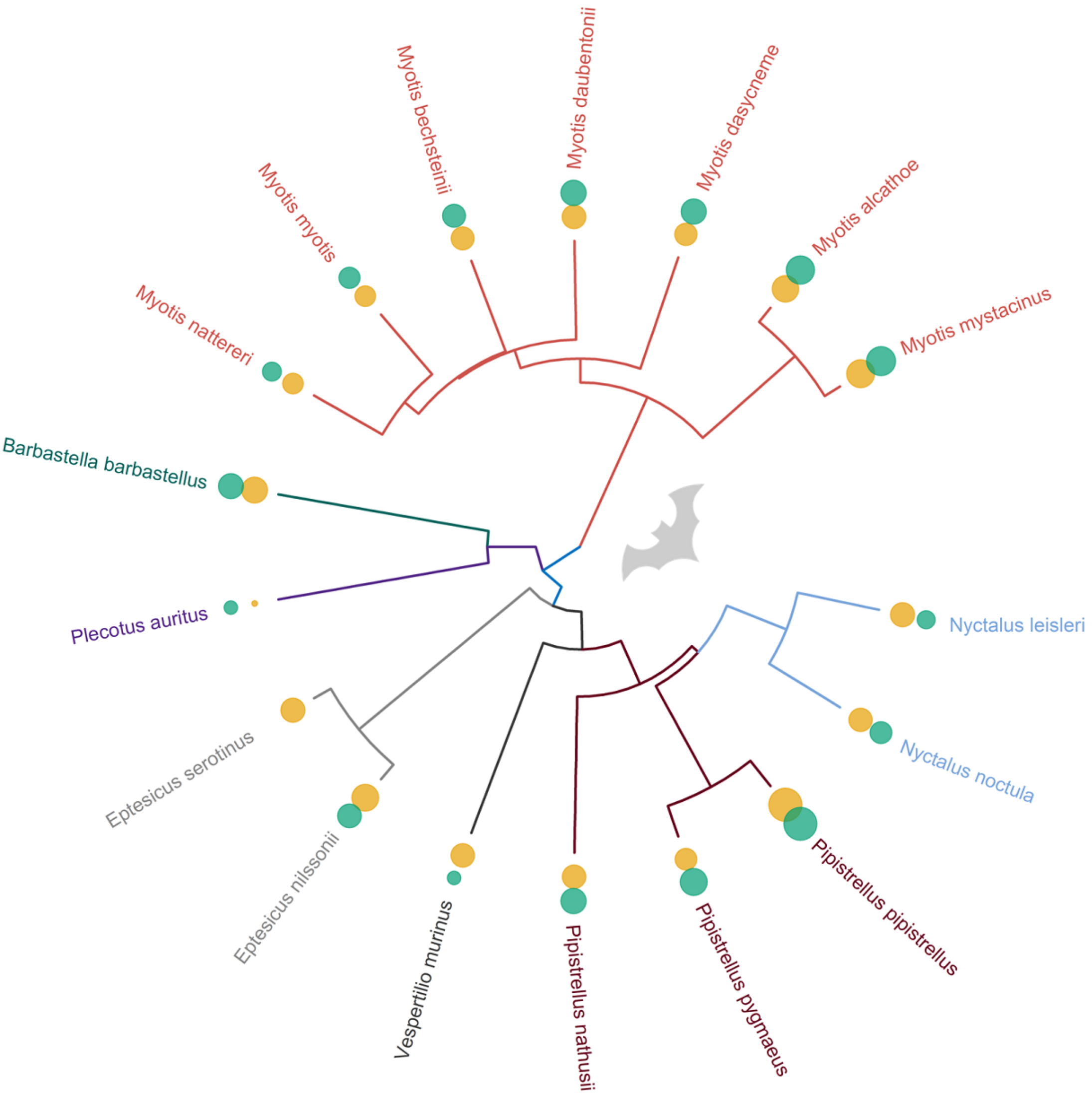


**Figure S1** Phylogenetic tree of all identified bat species across the eleven experimental forest landscapes. Branch colors indicate taxonomic genus. Dot sizes represent the $\log_e$-transformed total number of incidences per species in treatment districts (orange) and control districts (green).

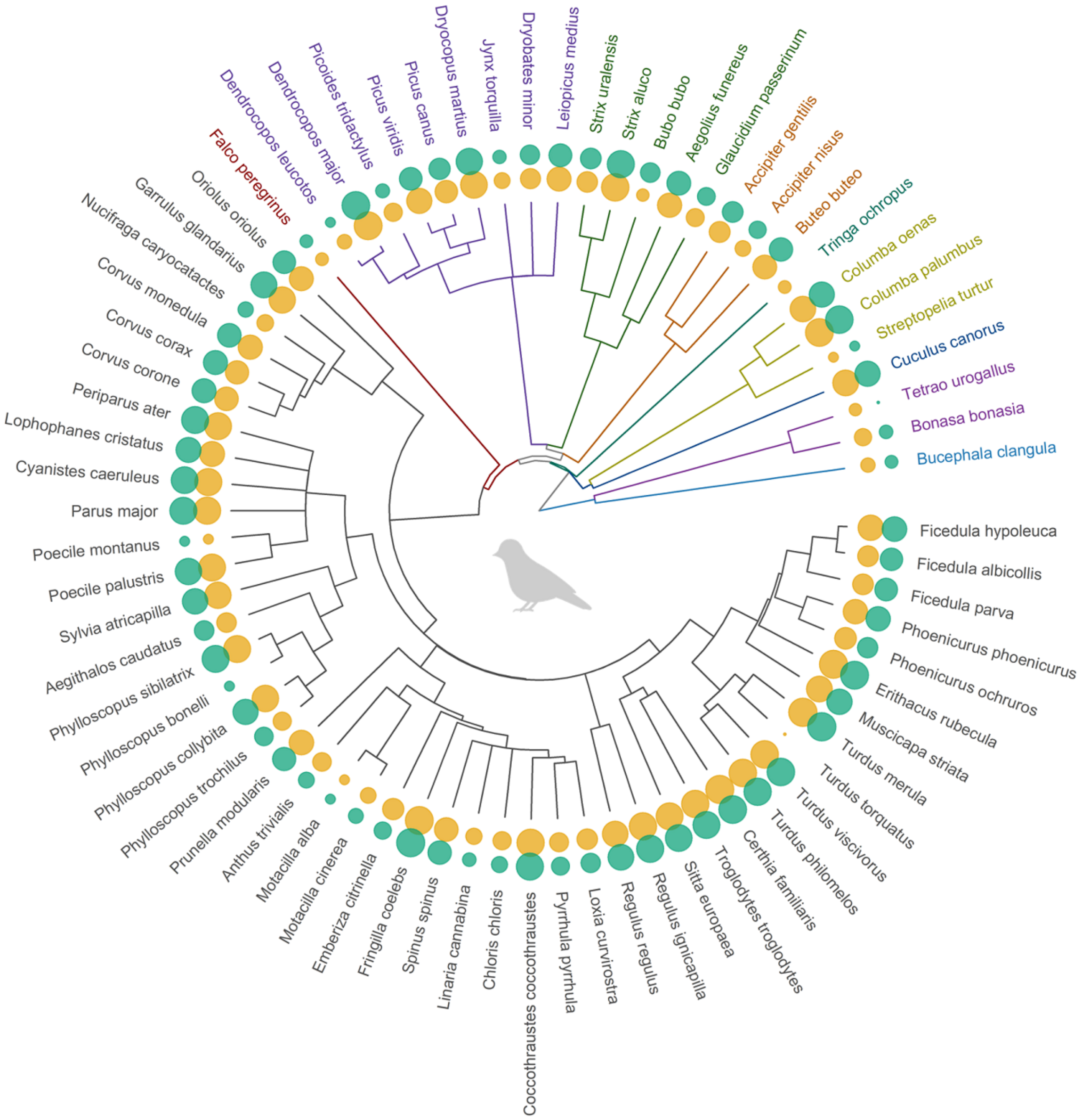


**Figure S2** Phylogenetic tree of all identified bird species across the eleven experimental forest landscapes. Branch colors indicate taxonomic orders. Dot sizes represent the $\log_e$-transformed total number of incidences per species in treatment districts (orange) and control districts (green).

**Table S1** Traits of identified bat species. All body size traits were log$_e$-transformed. Ear–Forearm and Tail–Forearm ratios represent residuals from linear models predicting ear and tail length from forearm length. Log$_e$-transformed Body Mass, Call Duration, and Maximum Frequency were standardized (mean = 0, SD = 1).

| Species | Aspect Ratio | Ear-Forearm | Tail-Forearm | Body Mass | Call Duration | Maximum Frequency | Guild |
|---|---|---|---|---|---|---|---|
| *Barbastella barbastellus* | 2.1 | -0.021 | 0.11 | -0.15 | -0.70 | -0.72 | edge |
| *Eptesicus nilssonii* | 2.1 | -0.0050 | 0.045 | 0.36 | 2.2 | -1.1 | open |
| *Eptesicus serotinus* | 2.2 | -0.15 | -0.0058 | 1.3 | 0.18 | -0.55 | open |
| *Myotis alcathoe* | 2.1 | 0.14 | 0.067 | -1.4 | -0.88 | 1.2 | edge |
| *Myotis bechsteinii* | 2.0 | 0.40 | 0.020 | -0.22 | -0.72 | 1.8 | closed |
| *Myotis dasycneme* | 2.2 | -0.026 | 0.075 | 0.55 | 0.53 | -0.32 | edge |
| *Myotis daubentonii* | 2.1 | -0.16 | -0.019 | -0.30 | -0.76 | 0.84 | edge |
| *Myotis myotis* | 2.1 | 0.059 | -0.069 | 1.7 | -0.17 | -0.38 | edge |
| *Myotis mystacinus* | 2.1 | 0.086 | 0.012 | -1.2 | -0.85 | 1.5 | edge |
| *Myotis nattereri* | 2.0 | 0.058 | 0.064 | -0.42 | -0.87 | 0.58 | closed |
| *Nyctalus leisleri* | 2.5 | -0.20 | -0.078 | 0.62 | 0.43 | -1.3 | open |
| *Nyctalus noctula* | 2.8 | -0.14 | -0.0080 | 1.7 | 0.77 | -1.5 | open |
| *Pipistrellus nathusii* | 2.1 | -0.10 | 0.0043 | -0.50 | -0.089 | -0.14 | open |
| *Pipistrellus pipistrellus* | 2.2 | -0.07 | -0.13 | -1.2 | -0.42 | 0.44 | edge |
| *Pipistrellus pygmaeus* | 2.2 | -0.50 | -0.14 | -1.1 | -0.37 | 0.90 | edge |
| *Plecotus auritus* | 2.1 | 0.86 | 0.15 | -0.45 | -0.63 | -0.57 | closed |
| *Vespertilio murinus* | 2.3 | -0.22 | -0.094 | 0.61 | 2.4 | -0.70 | open |

Preprint not peer reviewed

**Table S2** Traits of identified bird species. All body size traits were log$_e$-transformed. Beak Length, Tarsus Length, and Tail Length represent residuals from linear models, corrected for log$_e$-transformed Body mass. Body mass was standardized (mean = 0, SD = 1) after log$_e$-transformation.

| **Species** | **Hand Wing Index** | **Habitat Density** | **Trophic Level** | **Body Mass** | **Beak Length** | **Tarsus Length** | **Tail Length** |
|---|---|---|---|---|---|---|---|
| *Accipiter gentilis* | 36 | 1 | Carnivore | 1.7 | -0.29 | 0.25 | 0.28 |
| *Accipiter nisus* | 36 | 1 | Carnivore | 0.85 | -0.45 | 0.4 | 0.23 |
| *Buteo buteo* | 38 | 3 | Carnivore | 1.6 | -0.14 | 0.41 | 0.20 |
| *Bucephala clangula* | 51 | 3 | Carnivore | 1.7 | -0.41 | -0.45 | -0.81 |
| *Tringa ochropus* | 46 | 3 | Carnivore | 0.15 | 0.76 | 0.22 | -0.40 |
| *Columba oenas* | 42 | 3 | Herbivore | 1 | -0.49 | -0.39 | -0.11 |
| *Columba palumbus* | 41 | 2 | Herbivore | 1.3 | -0.53 | -0.35 | 0.0081 |
| *Streptopelia turtur* | 44 | 2 | Herbivore | 0.53 | -0.64 | -0.34 | 0.032 |
| *Cuculus canorus* | 44 | 2 | Carnivore | 0.43 | 0.050 | -0.33 | 0.45 |
| *Falco peregrinus* | 52 | 3 | Carnivore | 1.6 | -0.32 | -0.11 | -0.20 |
| *Bonasa bonasia* | 33 | 1 | Herbivore | 1.3 | -0.65 | -0.21 | -0.23 |
| *Tetrao urogallus* | 33 | 2 | Herbivore | 2.4 | -0.24 | -0.15 | 0.021 |
| *Aegithalos caudatus* | 22 | 2 | Carnivore | -1.1 | -0.52 | 0.070 | 0.59 |
| *Certhia familiaris* | 19 | 1 | Omnivore | -1.1 | 0.53 | -0.027 | 0.20 |
| *Corvus corax* | 4 | 3 | Omnivore | 1.7 | 0.61 | 0.23 | 0.21 |
| *Corvus corone* | 37 | 3 | Carnivore | 1.4 | 0.43 | 0.24 | 0.11 |
| *Corvus monedula* | 39 | 3 | Omnivore | 0.92 | 0.15 | 0.15 | -0.036 |
| *Garrulus glandarius* | 19 | 1 | Omnivore | 0.65 | 0.2 | 0.17 | 0.29 |
| *Nucifraga caryocatactes* | 24 | 1 | Omnivore | 0.73 | 0.77 | 0.14 | 0.0046 |

| Species | Hand Wing Index | Habitat Density | Trophic Level | Body Mass | Beak Length | Tarsus Length | Tail Length |
|---|---|---|---|---|---|---|---|
| *Emberiza citrinella* | 26 | 3 | Herbivore | -0.39 | -0.23 | -0.085 | 0.047 |
| *Chloris chloris* | 32 | 2 | Herbivore | -0.47 | 0.13 | -0.15 | -0.18 |
| *Coccothraustes coccothraustes* | 33 | 1 | Herbivore | 0.011 | 0.32 | -0.17 | -0.38 |
| *Fringilla coelebs* | 28 | 2 | Omnivore | -0.52 | 0.038 | -0.095 | 0.011 |
| *Linaria cannabina* | 34 | 2 | Herbivore | -0.64 | -0.13 | -0.17 | -0.091 |
| *Loxia curvirostra* | 38 | 1 | Herbivore | -0.23 | 0.40 | -0.30 | -0.26 |
| *Pyrrhula pyrrhula* | 25 | 2 | Herbivore | -0.51 | -0.098 | -0.090 | 0.041 |
| *Spinus spinus* | 36 | 1 | Herbivore | -0.88 | 0.040 | -0.27 | -0.17 |
| *Anthus trivialis* | 29 | 2 | Omnivore | -0.53 | -0.16 | 0.057 | -0.078 |
| *Motacilla alba* | 31 | 3 | Carnivore | -0.52 | 0.027 | 0.18 | 0.29 |
| *Motacilla cinerea* | 35 | 3 | Carnivore | -0.72 | 0.23 | 0.016 | 0.46 |
| *Erithacus rubecula* | 2 | 1 | Carnivore | -0.70 | -0.21 | 0.31 | -0.051 |
| *Ficedula albicollis* | 32 | 1 | Carnivore | -0.91 | -0.26 | -0.011 | -0.054 |
| *Ficedula hypoleuca* | 3 | 1 | Carnivore | -0.86 | -0.22 | 0.043 | -0.069 |
| *Ficedula parva* | 25 | 1 | Carnivore | -1.0 | -0.15 | -0.024 | 0.015 |
| *Muscicapa striata* | 3 | 1 | Carnivore | -0.77 | 0.10 | -0.20 | 0.052 |
| *Phoenicurus ochruros* | 23 | 3 | Omnivore | -0.75 | -0.048 | 0.23 | 0.028 |
| *Phoenicurus phoenicurus* | 26 | 1 | Carnivore | -0.82 | -0.051 | 0.23 | 0.023 |
| *Oriolus oriolus* | 38 | 1 | Omnivore | 0.22 | 0.31 | -0.19 | -0.10 |
| *Cyanistes caeruleus* | 21 | 1 | Omnivore | -0.99 | -0.27 | -0.044 | -0.02 |
| *Lophophanes cristatus* | 2 | 1 | Omnivore | -1.0 | -0.078 | 0.076 | -0.017 |
| *Parus major* | 19 | 2 | Omnivore | -0.75 | -0.057 | -0.014 | 0.045 |
| *Periparus ater* | 22 | 1 | Omnivore | -1.1 | -0.13 | 0.00039 | -0.069 |

| Species | Hand Wing Index | Habitat Density | Trophic Level | Body Mass | Beak Length | Tarsus Length | Tail Length |
|---|---|---|---|---|---|---|---|
| *Poecile montanus* | 18 | 1 | Omnivore | -0.99 | -0.066 | 0.019 | 0.12 |
| *Poecile palustris* | 18 | 1 | Omnivore | -0.99 | -0.12 | 0.00062 | 0.044 |
| *Phylloscopus bonelli* | 23 | 2 | Carnivore | -1.1 | -0.055 | 0.2 | 0.018 |
| *Phylloscopus collybita* | 18 | 1 | Carnivore | -1.2 | -0.15 | 0.29 | 0.0049 |
| *Phylloscopus sibilatrix* | 31 | 1 | Carnivore | -1.1 | -0.071 | 0.18 | 0.02 |
| *Phylloscopus trochilus* | 26 | 1 | Carnivore | -1.1 | -0.07 | 0.26 | 0.067 |
| *Prunella modularis* | 21 | 2 | Omnivore | -0.62 | -0.14 | 0.086 | -0.043 |
| *Regulus ignicapilla* | 22 | 1 | Carnivore | -1.4 | -0.04 | 0.22 | -0.091 |
| *Regulus regulus* | 22 | 1 | Carnivore | -1.4 | -0.086 | 0.13 | -0.088 |
| *Sitta europaea* | 24 | 1 | Carnivore | -0.62 | 0.41 | 0.073 | -0.29 |
| *Sylvia atricapilla* | 27 | 1 | Omnivore | -0.74 | -0.17 | 0.16 | 0.054 |
| *Troglodytes troglodytes* | 16 | 1 | Carnivore | -1.1 | 0.1 | 0.14 | -0.5 |
| *Turdus merula* | 23 | 2 | Omnivore | 0.38 | 0.079 | 0.14 | 0.072 |
| *Turdus philomelos* | 31 | 1 | Omnivore | 0.12 | -0.066 | 0.20 | -0.072 |
| *Turdus torquatus* | 32 | 2 | Omnivore | 0.41 | -0.047 | 0.099 | 0.086 |
| *Turdus viscivorus* | 35 | 2 | Omnivore | 0.46 | -0.025 | 0.076 | 0.084 |
| *Dendrocopos leucotos* | 3 | 1 | Carnivore | 0.46 | 0.64 | -0.20 | -0.11 |
| *Dendrocopos major* | 29 | 1 | Omnivore | 0.18 | 0.51 | -0.11 | -0.027 |
| *Dryobates minor* | 27 | 1 | Carnivore | -0.63 | 0.34 | -0.26 | -0.029 |
| *Dryocopus martius* | 26 | 1 | Carnivore | 1.1 | 0.86 | -0.11 | 0.25 |
| *Jynx torquilla* | 27 | 1 | Carnivore | -0.28 | -0.15 | -0.10 | -0.065 |
| *Leiopicus medius* | 28 | 1 | Carnivore | 0.037 | 0.29 | -0.25 | 0.0065 |

| Species | Hand Wing Index | Habitat Density | Trophic Level | Body Mass | Beak Length | Tarsus Length | Tail Length |
|---|---|---|---|---|---|---|---|
| *Picoides tridactylus* | 29 | 1 | Carnivore | 0.10 | 0.58 | -0.27 | -0.068 |
| *Picus canus* | 26 | 1 | Carnivore | 0.56 | 0.47 | -0.24 | 0.0063 |
| *Picus viridis* | 25 | 2 | Carnivore | 0.71 | 0.74 | -0.15 | -0.11 |
| *Aegolius funereus* | 32 | 1 | Carnivore | 0.56 | -0.35 | -0.30 | -0.047 |
| *Bubo bubo* | 33 | 2 | Carnivore | 2.4 | -0.31 | 0.11 | -0.024 |
| *Glaucidium passerinum* | 26 | 1 | Carnivore | 0.025 | -0.33 | -0.48 | -0.36 |
| *Strix aluco* | 29 | 1 | Carnivore | 1.3 | -0.31 | 0.045 | 0.11 |
| *Strix uralensis* | 29 | 2 | Carnivore | 1.6 | -0.26 | -0.002 | 0.39 |

Preprint not peer reviewed

**Gamma with Coverage = 0.819**

| Site | TD(q=0) E | TD(q=0) C | Difference | LCL | UCL | W(fixed) |
|---|---|---|---|---|---|---|
| B04 | 11.651 | 11.158 | 0.493 | -4.938 | 5.923 | 9.91% |
| B05 | 10.328 | 6.777 | 3.551 | -1.573 | 8.674 | 11.13% |
| B06 | 9.869 | 8.471 | 1.398 | -1.009 | 3.806 | 50.4% |
| B07 | 13.102 | 5.307 | 7.795 | -0.731 | 16.321 | 4.02% |
| H09 | 21.106 | 15.92 | 5.187 | -12.552 | 22.926 | 0.93% |
| L11 | 6.784 | 12.21 | -5.426 | -12.52 | 1.668 | 5.81% |
| P08 | 15.104 | 11.741 | 3.363 | -8.4 | 15.126 | 2.11% |
| S10 | 16.513 | 3.553 | 12.96 | -0.3 | 26.221 | 1.66% |
| U01 | 11.999 | 9.384 | 2.616 | -4.699 | 9.931 | 5.46% |
| U02 | 14.637 | 11.05 | 3.587 | -6.479 | 13.653 | 2.88% |
| U03 | 13.544 | 13.148 | 0.396 | -6.766 | 7.558 | 5.7% |
| **Meta analysis** | | | **1.75** | **0.041** | **3.46** | |

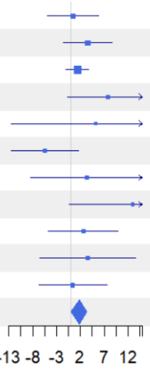


Figure S3

**Gamma with Coverage = 0.819**

| Site | TD(q=1) E | TD(q=1) C | Difference | LCL | UCL | W(fixed) |
|---|---|---|---|---|---|---|
| B04 | 11.081 | 9.926 | 1.155 | -2.316 | 4.625 | 12% |
| B05 | 9.538 | 6.318 | 3.219 | 0.434 | 6.005 | 18.63% |
| B06 | 9.553 | 7.956 | 1.597 | -0.324 | 3.518 | 39.18% |
| B07 | 11.513 | 5.307 | 6.206 | 2.13 | 10.282 | 8.7% |
| H09 | 16.427 | 13.535 | 2.892 | -12.662 | 18.447 | 0.6% |
| L11 | 6.429 | 11.57 | -5.142 | -12.396 | 2.112 | 2.75% |
| P08 | 12.204 | 10.564 | 1.641 | -5.916 | 9.197 | 2.53% |
| S10 | 13.439 | 3.324 | 10.116 | -0.283 | 20.514 | 1.34% |
| U01 | 11.046 | 8.787 | 2.259 | -2.393 | 6.911 | 6.68% |
| U02 | 12.845 | 10.402 | 2.443 | -3.731 | 8.618 | 3.79% |
| U03 | 12.314 | 11.759 | 0.555 | -5.621 | 6.731 | 3.79% |
| **Meta analysis** | | | **2.222** | **1.02** | **3.424** | |

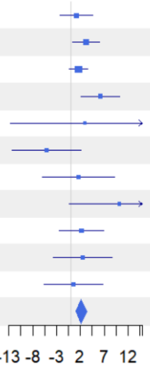


Figure S4

**Gamma with Coverage = 0.819**

| Site | TD(q=2) E | TD(q=2) C | Difference | LCL | UCL | W(fixed) |
|---|---|---|---|---|---|---|
| B04 | 10.608 | 9.002 | 1.606 | -1.098 | 4.309 | 11.39% |
| B05 | 8.913 | 5.954 | 2.959 | 0.978 | 4.94 | 21.22% |
| B06 | 9.303 | 7.525 | 1.778 | -0.123 | 3.678 | 23.04% |
| B07 | 10.33 | 5.307 | 5.024 | 2.581 | 7.466 | 13.96% |
| H09 | 12.101 | 10.506 | 1.596 | -4.926 | 8.117 | 1.96% |
| L11 | 6.134 | 11.036 | -4.901 | -9.048 | -0.754 | 4.84% |
| P08 | 10.108 | 9.795 | 0.313 | -4.492 | 5.119 | 3.61% |
| S10 | 9.205 | 3.121 | 6.084 | 0.508 | 11.661 | 2.68% |
| U01 | 10.292 | 8.28 | 2.012 | -1.657 | 5.681 | 6.18% |
| U02 | 11.337 | 9.811 | 1.526 | -2.221 | 5.273 | 5.93% |
| U03 | 11.368 | 10.745 | 0.623 | -3.38 | 4.626 | 5.2% |
| **Meta analysis** | | | **2.137** | **1.225** | **3.049** | |

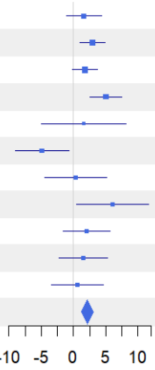


Figure S5

Preprint not peer reviewed

**Gamma with Coverage = 0.819**

| Site | PD(q=0) E | PD(q=0) C | Difference | LCL | UCL | W(fixed) |
|---|---|---|---|---|---|---|
| B04 | 6.024 | 6.082 | -0.058 | -3.482 | 3.366 | 7.47% |
| B05 | 5.315 | 3.683 | 1.632 | -2.126 | 5.391 | 6.2% |
| B06 | 5.684 | 4.813 | 0.871 | -3.335 | 5.077 | 4.95% |
| B07 | 6.988 | 3.214 | 3.774 | 1.184 | 6.364 | 13.05% |
| H09 | 10.522 | 7 | 3.522 | -1.07 | 8.113 | 4.15% |
| L11 | 3.545 | 6.649 | -3.105 | -7.371 | 1.162 | 4.81% |
| P08 | 7.307 | 6.714 | 0.594 | -3.76 | 4.947 | 4.62% |
| S10 | 7.121 | 2.518 | 4.604 | 0.957 | 8.25 | 6.58% |
| U01 | 6.046 | 5.056 | 0.99 | -0.98 | 2.96 | 22.56% |
| U02 | 7.268 | 5.66 | 1.608 | -2.438 | 5.654 | 5.35% |
| U03 | 6.61 | 6.07 | 0.54 | -1.539 | 2.619 | 20.25% |
| **Meta analysis** | | | **1.379** | **0.443** | **2.314** | |

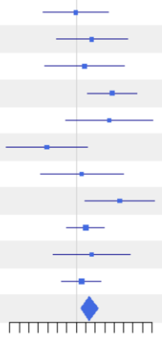


Figure S6

**Gamma with Coverage = 0.819**

| Site | PD(q=1) E | PD(q=1) C | Difference | LCL | UCL | W(fixed) |
|---|---|---|---|---|---|---|
| B04 | 5.771 | 5.494 | 0.277 | -2.565 | 3.118 | 6.21% |
| B05 | 4.963 | 3.486 | 1.477 | -1.062 | 4.017 | 7.77% |
| B06 | 5.528 | 4.546 | 0.982 | -2.321 | 4.286 | 4.59% |
| B07 | 6.346 | 3.214 | 3.131 | 1.093 | 5.17 | 12.07% |
| H09 | 8.064 | 5.489 | 2.575 | -0.527 | 5.677 | 5.21% |
| L11 | 3.391 | 6.351 | -2.96 | -5.809 | -0.111 | 6.18% |
| P08 | 6.139 | 6.165 | -0.026 | -3.147 | 3.094 | 5.15% |
| S10 | 6.027 | 2.406 | 3.621 | 1.28 | 5.962 | 9.15% |
| U01 | 5.621 | 4.752 | 0.869 | -0.844 | 2.583 | 17.08% |
| U02 | 6.324 | 5.33 | 0.994 | -2.676 | 4.663 | 3.72% |
| U03 | 6.009 | 5.553 | 0.456 | -1.024 | 1.937 | 22.87% |
| **Meta analysis** | | | **1.126** | **0.418** | **1.834** | |

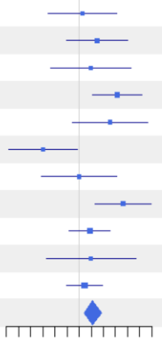


Figure S7

**Gamma with Coverage = 0.819**

| Site | PD(q=2) E | PD(q=2) C | Difference | LCL | UCL | W(fixed) |
|---|---|---|---|---|---|---|
| B04 | 5.578 | 5.11 | 0.468 | -1.943 | 2.878 | 5.94% |
| B05 | 4.715 | 3.352 | 1.363 | -0.647 | 3.372 | 8.55% |
| B06 | 5.412 | 4.345 | 1.067 | -1.702 | 3.836 | 4.5% |
| B07 | 5.938 | 3.214 | 2.724 | 0.898 | 4.55 | 10.36% |
| H09 | 6.406 | 4.447 | 1.959 | -0.467 | 4.385 | 5.87% |
| L11 | 3.271 | 6.085 | -2.814 | -4.943 | -0.685 | 7.62% |
| P08 | 5.506 | 5.833 | -0.327 | -2.719 | 2.065 | 6.04% |
| S10 | 4.887 | 2.313 | 2.574 | 0.951 | 4.197 | 13.1% |
| U01 | 5.303 | 4.499 | 0.804 | -0.784 | 2.392 | 13.69% |
| U02 | 5.683 | 5.038 | 0.645 | -2.377 | 3.666 | 3.78% |
| U03 | 5.574 | 5.224 | 0.349 | -0.947 | 1.646 | 20.54% |
| **Meta analysis** | | | **0.899** | **0.311** | **1.487** | |

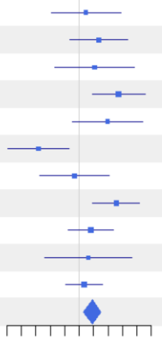


Figure S8

Gamma with Coverage = 0.762

| Site | FD(q=0) E | FD(q=0) C | Difference | LCL | UCL | W(fixed) |
|---|---|---|---|---|---|---|
| B04 | 3.683 | 2.728 | 0.956 | -0.018 | 1.93 | 13.47% |
| B05 | 2.589 | 1.957 | 0.632 | -0.623 | 1.887 | 8.11% |
| B06 | 3.175 | 2.547 | 0.628 | -0.459 | 1.715 | 10.82% |
| B07 | 3.323 | 2.184 | 1.139 | -0.05 | 2.328 | 9.04% |
| H09 | 3.043 | 3.219 | -0.176 | -1.446 | 1.094 | 7.92% |
| L11 | 2.507 | 2.823 | -0.316 | -1.841 | 1.209 | 5.5% |
| P08 | 3.462 | 3.719 | -0.257 | -1.772 | 1.259 | 5.57% |
| S10 | 2.698 | 1.436 | 1.262 | -0.486 | 3.009 | 4.19% |
| U01 | 2.784 | 2.189 | 0.596 | -0.338 | 1.529 | 14.67% |
| U02 | 2.737 | 2.928 | -0.191 | -1.633 | 1.251 | 6.14% |
| U03 | 2.978 | 2.45 | 0.528 | -0.409 | 1.464 | 14.57% |
| **Meta analysis** | | | **0.511** | **0.153** | **0.868** | |

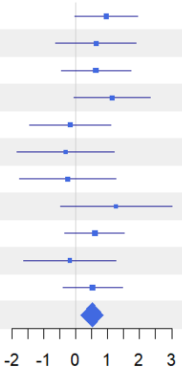


Figure S9

Gamma with Coverage = 0.762

| Site | FD(q=1) E | FD(q=1) C | Difference | LCL | UCL | W(fixed) |
|---|---|---|---|---|---|---|
| B04 | 3.091 | 2.21 | 0.881 | 0.159 | 1.603 | 11.55% |
| B05 | 2.157 | 1.769 | 0.388 | -0.335 | 1.11 | 11.53% |
| B06 | 2.625 | 2.128 | 0.497 | -0.293 | 1.287 | 9.66% |
| B07 | 2.841 | 1.985 | 0.856 | 0.053 | 1.659 | 9.34% |
| H09 | 2.43 | 2.508 | -0.078 | -1.212 | 1.055 | 4.69% |
| L11 | 2.179 | 2.389 | -0.21 | -1.356 | 0.935 | 4.59% |
| P08 | 2.863 | 3.283 | -0.42 | -1.513 | 0.674 | 5.04% |
| S10 | 2.038 | 1.34 | 0.698 | -0.33 | 1.726 | 5.69% |
| U01 | 2.336 | 1.953 | 0.383 | -0.236 | 1.002 | 15.72% |
| U02 | 2.213 | 2.515 | -0.302 | -1.154 | 0.55 | 8.3% |
| U03 | 2.466 | 2.105 | 0.361 | -0.297 | 1.019 | 13.91% |
| **Meta analysis** | | | **0.365** | **0.12** | **0.61** | |

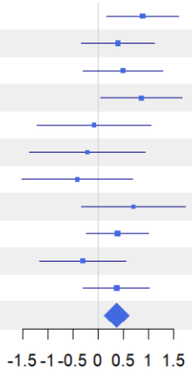


Figure S10

Gamma with Coverage = 0.762

| Site | FD(q=2) E | FD(q=2) C | Difference | LCL | UCL | W(fixed) |
|---|---|---|---|---|---|---|
| B04 | 2.673 | 1.919 | 0.753 | 0.186 | 1.321 | 11.41% |
| B05 | 1.902 | 1.638 | 0.264 | -0.293 | 0.822 | 11.83% |
| B06 | 2.288 | 1.889 | 0.399 | -0.298 | 1.097 | 7.56% |
| B07 | 2.502 | 1.837 | 0.665 | 0.066 | 1.265 | 10.23% |
| H09 | 2.073 | 2.103 | -0.03 | -0.831 | 0.771 | 5.72% |
| L11 | 1.957 | 2.118 | -0.161 | -1.028 | 0.706 | 4.89% |
| P08 | 2.488 | 2.956 | -0.468 | -1.54 | 0.605 | 3.2% |
| S10 | 1.724 | 1.269 | 0.455 | -0.129 | 1.04 | 10.77% |
| U01 | 2.062 | 1.793 | 0.269 | -0.274 | 0.813 | 12.44% |
| U02 | 1.926 | 2.247 | -0.321 | -1.105 | 0.464 | 5.97% |
| U03 | 2.165 | 1.894 | 0.271 | -0.209 | 0.75 | 16% |
| **Meta analysis** | | | **0.298** | **0.106** | **0.489** | |

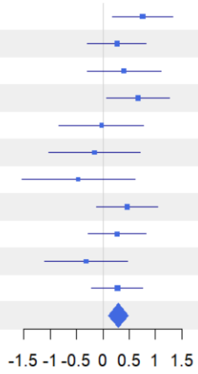


Figure S11

**1-S with Coverage = 0.819**

| Site | TD(q=0) E | TD(q=0) C | Difference | LCL | UCL | W(fixed) |
|---|---|---|---|---|---|---|
| B04 | 0.478 | 0.493 | -0.015 | -0.205 | 0.174 | 9.68% |
| B05 | 0.499 | 0.482 | 0.017 | -0.144 | 0.179 | 13.31% |
| B06 | 0.172 | 0.54 | -0.368 | -0.679 | -0.057 | 3.58% |
| B07 | 0.588 | 0.273 | 0.315 | 0.053 | 0.577 | 5.05% |
| H09 | 0.717 | -1.407 | 2.123 | 1.893 | 2.354 | 6.51% |
| L11 | 0.499 | 0.709 | -0.21 | -0.421 | 0 | 7.82% |
| P08 | 0.708 | 0.243 | 0.465 | 0.079 | 0.851 | 2.32% |
| S10 | 0.967 | 0.278 | 0.689 | 0.255 | 1.122 | 1.84% |
| U01 | 0.59 | 0.466 | 0.124 | -0.007 | 0.255 | 20.15% |
| U02 | 0.735 | 0.245 | 0.489 | 0.298 | 0.681 | 9.46% |
| U03 | 0.618 | 0.686 | -0.068 | -0.199 | 0.063 | 20.28% |
| **Meta analysis** | | | **0.206** | **0.148** | **0.265** | |

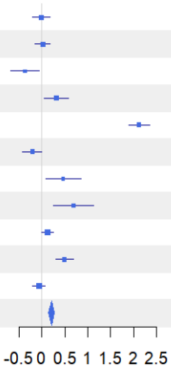


Figure S12

**1-S with Coverage = 0.819**

| Site | TD(q=1) E | TD(q=1) C | Difference | LCL | UCL | W(fixed) |
|---|---|---|---|---|---|---|
| B04 | 0.538 | 0.576 | -0.038 | -0.192 | 0.116 | 7.58% |
| B05 | 0.58 | 0.518 | 0.062 | -0.075 | 0.2 | 9.48% |
| B06 | 0.363 | 0.576 | -0.213 | -0.42 | -0.006 | 4.18% |
| B07 | 0.591 | 0.38 | 0.211 | 0.037 | 0.385 | 5.91% |
| H09 | 0.716 | -0.393 | 1.109 | 0.926 | 1.293 | 5.33% |
| L11 | 0.489 | 0.707 | -0.218 | -0.409 | -0.027 | 4.91% |
| P08 | 0.676 | 0.25 | 0.426 | 0.234 | 0.618 | 4.84% |
| S10 | 0.939 | 0.341 | 0.598 | 0.344 | 0.851 | 2.79% |
| U01 | 0.607 | 0.502 | 0.105 | 0.022 | 0.188 | 25.81% |
| U02 | 0.732 | 0.272 | 0.46 | 0.338 | 0.582 | 12.01% |
| U03 | 0.625 | 0.684 | -0.059 | -0.161 | 0.043 | 17.17% |
| **Meta analysis** | | | **0.165** | **0.122** | **0.207** | |

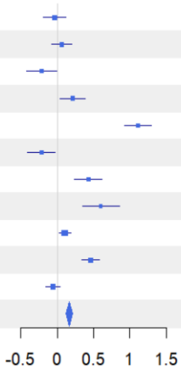


Figure S13

**1-S with Coverage = 0.819**

| Site | TD(q=2) E | TD(q=2) C | Difference | LCL | UCL | W(fixed) |
|---|---|---|---|---|---|---|
| B04 | 0.596 | 0.665 | -0.069 | -0.158 | 0.02 | 9.64% |
| B05 | 0.655 | 0.545 | 0.11 | 0.011 | 0.208 | 7.91% |
| B06 | 0.542 | 0.632 | -0.089 | -0.192 | 0.013 | 7.34% |
| B07 | 0.606 | 0.478 | 0.128 | 0.023 | 0.233 | 6.98% |
| H09 | 0.727 | 0.604 | 0.123 | 0.014 | 0.231 | 6.51% |
| L11 | 0.502 | 0.709 | -0.207 | -0.357 | -0.056 | 3.4% |
| P08 | 0.698 | 0.392 | 0.306 | 0.189 | 0.424 | 5.59% |
| S10 | 0.868 | 0.45 | 0.418 | 0.249 | 0.587 | 2.69% |
| U01 | 0.635 | 0.534 | 0.1 | 0.047 | 0.154 | 26.57% |
| U02 | 0.755 | 0.338 | 0.416 | 0.32 | 0.512 | 8.25% |
| U03 | 0.647 | 0.687 | -0.04 | -0.111 | 0.031 | 15.12% |
| **Meta analysis** | | | **0.089** | **0.061** | **0.116** | |

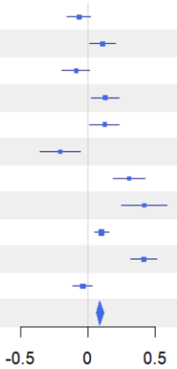


Figure S14

**1-S with Coverage = 0.819**

| Site | PD(q=0) E | PD(q=0) C | Difference | LCL | UCL | W(fixed) |
|---|---|---|---|---|---|---|
| B04 | 0.312 | 0.418 | -0.106 | -0.418 | 0.206 | 5.41% |
| B05 | 0.37 | 0.321 | 0.05 | -0.257 | 0.357 | 5.59% |
| B06 | 0.221 | 0.414 | -0.193 | -0.727 | 0.34 | 1.85% |
| B07 | 0.376 | 0.146 | 0.23 | -0.093 | 0.553 | 5.04% |
| H09 | 0.641 | -0.374 | 1.015 | 0.706 | 1.324 | 5.53% |
| L11 | 0.334 | 0.667 | -0.333 | -0.579 | -0.087 | 8.71% |
| P08 | 0.494 | 0.094 | 0.4 | 0.059 | 0.74 | 4.55% |
| S10 | 0.797 | 0.027 | 0.77 | 0.457 | 1.083 | 5.37% |
| U01 | 0.501 | 0.405 | 0.096 | -0.057 | 0.25 | 22.45% |
| U02 | 0.678 | 0.207 | 0.471 | 0.078 | 0.864 | 3.41% |
| U03 | 0.492 | 0.555 | -0.063 | -0.191 | 0.065 | 32.07% |
| **Meta analysis** | | | **0.109** | **0.037** | **0.182** | |

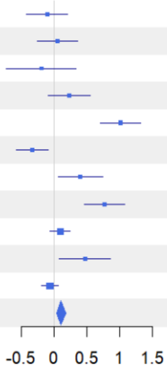


Figure S15

**1-S with Coverage = 0.819**

| Site | PD(q=1) E | PD(q=1) C | Difference | LCL | UCL | W(fixed) |
|---|---|---|---|---|---|---|
| B04 | 0.394 | 0.477 | -0.083 | -0.326 | 0.161 | 4.78% |
| B05 | 0.44 | 0.356 | 0.084 | -0.146 | 0.313 | 5.38% |
| B06 | 0.355 | 0.467 | -0.112 | -0.54 | 0.316 | 1.55% |
| B07 | 0.405 | 0.27 | 0.135 | -0.134 | 0.404 | 3.92% |
| H09 | 0.606 | 0.061 | 0.546 | 0.369 | 0.722 | 9.12% |
| L11 | 0.328 | 0.669 | -0.341 | -0.532 | -0.149 | 7.74% |
| P08 | 0.481 | 0.165 | 0.316 | 0.046 | 0.587 | 3.87% |
| S10 | 0.754 | 0.152 | 0.602 | 0.347 | 0.857 | 4.35% |
| U01 | 0.52 | 0.429 | 0.091 | -0.026 | 0.208 | 20.82% |
| U02 | 0.654 | 0.241 | 0.413 | 0.098 | 0.729 | 2.85% |
| U03 | 0.5 | 0.546 | -0.046 | -0.135 | 0.044 | 35.61% |
| **Meta analysis** | | | **0.08** | **0.027** | **0.134** | |

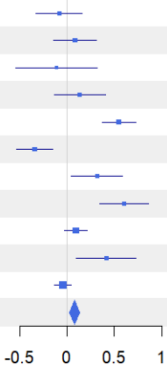


Figure S16

**1-S with Coverage = 0.819**

| Site | PD(q=2) E | PD(q=2) C | Difference | LCL | UCL | W(fixed) |
|---|---|---|---|---|---|---|
| B04 | 0.456 | 0.546 | -0.09 | -0.306 | 0.127 | 3.96% |
| B05 | 0.497 | 0.382 | 0.115 | -0.073 | 0.303 | 5.24% |
| B06 | 0.474 | 0.534 | -0.06 | -0.409 | 0.289 | 1.52% |
| B07 | 0.446 | 0.373 | 0.073 | -0.16 | 0.306 | 3.42% |
| H09 | 0.618 | 0.41 | 0.208 | 0.093 | 0.323 | 14.04% |
| L11 | 0.339 | 0.672 | -0.333 | -0.487 | -0.18 | 7.82% |
| P08 | 0.528 | 0.268 | 0.259 | 0.03 | 0.488 | 3.53% |
| S10 | 0.688 | 0.276 | 0.412 | 0.198 | 0.626 | 4.04% |
| U01 | 0.541 | 0.443 | 0.098 | -0.006 | 0.202 | 17.03% |
| U02 | 0.657 | 0.292 | 0.365 | 0.105 | 0.625 | 2.74% |
| U03 | 0.515 | 0.544 | -0.029 | -0.1 | 0.042 | 36.67% |
| **Meta analysis** | | | **0.049** | **0.006** | **0.092** | |

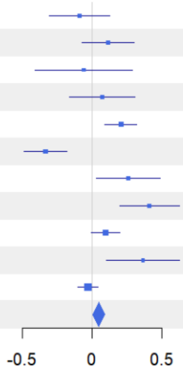


Figure S17

1-S with Coverage = 0.762

| Site | FD(q=0) E | FD(q=0) C | | Difference | LCL | UCL | W(fixed) |
|---|---|---|---|---|---|---|---|
| B04 | -0.006 | -0.164 | | 0.158 | -0.077 | 0.393 | 14.77% |
| B05 | -0.083 | -0.045 | | -0.038 | -0.333 | 0.258 | 9.36% |
| B06 | -0.175 | -0.118 | | -0.057 | -0.395 | 0.281 | 7.15% |
| B07 | -0.064 | -0.048 | | -0.015 | -0.365 | 0.334 | 6.69% |
| H09 | -0.132 | -0.428 | | 0.295 | -0.238 | 0.829 | 2.87% |
| L11 | -0.1 | -0.06 | | -0.04 | -0.425 | 0.345 | 5.51% |
| P08 | -0.053 | -0.277 | | 0.224 | -0.134 | 0.581 | 6.4% |
| S10 | 0.189 | -0.26 | | 0.449 | -0.095 | 0.993 | 2.76% |
| U01 | -0.064 | -0.073 | | 0.01 | -0.221 | 0.241 | 15.35% |
| U02 | -0.159 | -0.146 | | -0.013 | -0.334 | 0.308 | 7.94% |
| U03 | -0.084 | -0.047 | | -0.037 | -0.233 | 0.159 | 21.18% |
| **Meta analysis** | | | | **0.04** | **-0.05** | **0.131** | |

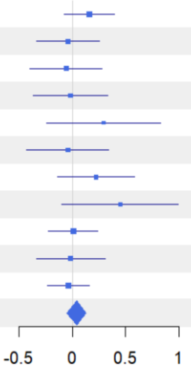


Figure S18

1-S with Coverage = 0.762

| Site | FD(q=1) E | FD(q=1) C | | Difference | LCL | UCL | W(fixed) |
|---|---|---|---|---|---|---|---|
| B04 | -0.003 | -0.14 | | 0.137 | -0.11 | 0.384 | 9.88% |
| B05 | -0.08 | 0.024 | | -0.103 | -0.308 | 0.102 | 14.31% |
| B06 | -0.092 | -0.026 | | -0.066 | -0.307 | 0.174 | 10.36% |
| B07 | -0.078 | 0.041 | | -0.119 | -0.373 | 0.135 | 9.32% |
| H09 | -0.191 | -0.428 | | 0.238 | -0.121 | 0.596 | 4.67% |
| L11 | -0.068 | -0.082 | | 0.014 | -0.405 | 0.432 | 3.43% |
| P08 | 0.004 | -0.267 | | 0.27 | -0.122 | 0.663 | 3.9% |
| S10 | 0.011 | -0.309 | | 0.32 | -0.199 | 0.839 | 2.23% |
| U01 | -0.065 | -0.038 | | -0.027 | -0.204 | 0.15 | 19.14% |
| U02 | -0.031 | -0.155 | | 0.124 | -0.121 | 0.369 | 10.03% |
| U03 | -0.094 | -0.014 | | -0.08 | -0.297 | 0.138 | 12.73% |
| **Meta analysis** | | | | **0.007** | **-0.07** | **0.085** | |

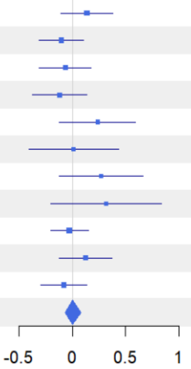


Figure S19

1-S with Coverage = 0.762

| Site | FD(q=2) E | FD(q=2) C | | Difference | LCL | UCL | W(fixed) |
|---|---|---|---|---|---|---|---|
| B04 | -0.009 | -0.105 | | 0.096 | -0.1 | 0.292 | 9.66% |
| B05 | -0.064 | 0.071 | | -0.135 | -0.289 | 0.019 | 15.57% |
| B06 | -0.025 | 0.071 | | -0.097 | -0.348 | 0.155 | 5.87% |
| B07 | -0.083 | 0.131 | | -0.214 | -0.414 | -0.013 | 9.25% |
| H09 | -0.204 | -0.298 | | 0.095 | -0.185 | 0.374 | 4.76% |
| L11 | -0.022 | -0.078 | | 0.057 | -0.24 | 0.353 | 4.23% |
| P08 | 0.14 | -0.149 | | 0.288 | 0.015 | 0.562 | 4.96% |
| S10 | -0.056 | -0.258 | | 0.202 | -0.152 | 0.555 | 2.97% |
| U01 | -0.047 | 0.001 | | -0.048 | -0.199 | 0.103 | 16.27% |
| U02 | 0.124 | -0.143 | | 0.267 | 0.073 | 0.46 | 9.96% |
| U03 | -0.078 | 0.025 | | -0.103 | -0.253 | 0.047 | 16.5% |
| **Meta analysis** | | | | **-0.008** | **-0.069** | **0.053** | |

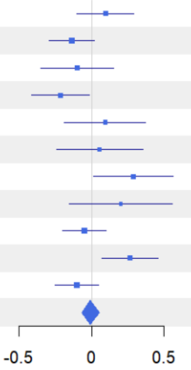


Figure S20

**Alpha with Coverage = 0.819**

| Site | TD(q=0) E | TD(q=0) C | Difference | LCL | UCL | W(fixed) |
|---|---|---|---|---|---|---|
| B04 | 6.702 | 6.266 | 0.436 | -2.245 | 3.116 | 6.51% |
| B05 | 5.744 | 3.874 | 1.869 | -1.478 | 5.217 | 4.18% |
| B06 | 8.358 | 4.405 | 3.953 | 0.102 | 7.804 | 3.16% |
| B07 | 6.253 | 4.037 | 2.216 | -0.02 | 4.453 | 9.36% |
| H09 | 7.663 | 35.828 | -28.165 | -57.424 | 1.093 | 0.05% |
| L11 | 3.775 | 4.513 | -0.738 | -2.675 | 1.199 | 12.48% |
| P08 | 5.751 | 9.206 | -3.454 | -8.455 | 1.546 | 1.87% |
| S10 | 2.325 | 2.689 | -0.365 | -2.151 | 1.422 | 14.67% |
| U01 | 5.395 | 5.305 | 0.089 | -1.635 | 1.814 | 15.73% |
| U02 | 4.652 | 8.534 | -3.881 | -6.242 | -1.521 | 8.4% |
| U03 | 5.735 | 4.734 | 1.001 | -0.407 | 2.41 | 23.59% |
| **Meta analysis** | | | **0.137** | **-0.547** | **0.821** | |

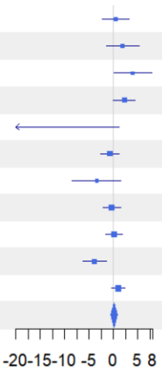


Figure S21

**Alpha with Coverage = 0.819**

| Site | TD(q=1) E | TD(q=1) C | Difference | LCL | UCL | W(fixed) |
|---|---|---|---|---|---|---|
| B04 | 5.783 | 4.847 | 0.937 | -0.839 | 2.712 | 6.15% |
| B05 | 4.619 | 3.409 | 1.21 | 0.13 | 2.289 | 16.63% |
| B06 | 6.47 | 3.881 | 2.589 | 0.514 | 4.664 | 4.5% |
| B07 | 5.468 | 3.544 | 1.924 | 0.706 | 3.141 | 13.07% |
| H09 | 5.968 | 18.263 | -12.295 | -22.682 | -1.908 | 0.18% |
| L11 | 3.635 | 4.295 | -0.661 | -1.922 | 0.6 | 12.19% |
| P08 | 4.989 | 8.218 | -3.23 | -6.517 | 0.058 | 1.79% |
| S10 | 2.218 | 2.331 | -0.112 | -1.804 | 1.579 | 6.78% |
| U01 | 4.79 | 4.673 | 0.117 | -0.946 | 1.181 | 17.13% |
| U02 | 4.109 | 7.772 | -3.663 | -5.44 | -1.886 | 6.14% |
| U03 | 5.131 | 4.253 | 0.879 | -0.242 | 2 | 15.42% |
| **Meta analysis** | | | **0.389** | **-0.051** | **0.83** | |

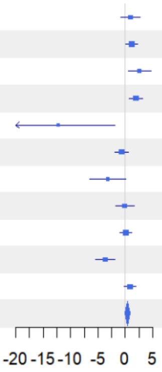


Figure S22

**Alpha with Coverage = 0.819**

| Site | TD(q=2) E | TD(q=2) C | Difference | LCL | UCL | W(fixed) |
|---|---|---|---|---|---|---|
| B04 | 4.984 | 3.678 | 1.306 | 0.331 | 2.281 | 9.01% |
| B05 | 3.723 | 3.068 | 0.655 | -0.045 | 1.356 | 17.44% |
| B06 | 4.82 | 3.3 | 1.52 | 0.463 | 2.576 | 7.67% |
| B07 | 4.763 | 3.086 | 1.677 | 0.814 | 2.539 | 11.52% |
| H09 | 4.284 | 4.864 | -0.58 | -2.758 | 1.598 | 1.81% |
| L11 | 3.397 | 4.084 | -0.687 | -1.975 | 0.601 | 5.16% |
| P08 | 3.933 | 6.384 | -2.451 | -4.08 | -0.822 | 3.23% |
| S10 | 2.104 | 1.893 | 0.211 | -0.937 | 1.358 | 6.5% |
| U01 | 4.196 | 4.151 | 0.045 | -0.829 | 0.919 | 11.21% |
| U02 | 3.393 | 6.727 | -3.334 | -4.541 | -2.127 | 5.88% |
| U03 | 4.509 | 3.859 | 0.649 | 0.004 | 1.294 | 20.58% |
| **Meta analysis** | | | **0.373** | **0.08** | **0.666** | |

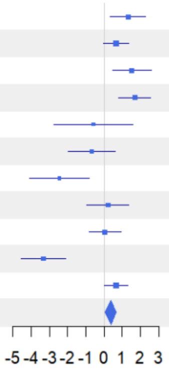


Figure S23

**Alpha with Coverage = 0.819**

| Site | PD(q=0) E | PD(q=0) C | Difference | LCL | UCL | W(fixed) |
|---|---|---|---|---|---|---|
| B04 | 4.352 | 3.82 | 0.532 | -0.944 | 2.007 | 4.31% |
| B05 | 3.565 | 2.633 | 0.932 | -0.034 | 1.898 | 10.05% |
| B06 | 4.568 | 3.042 | 1.527 | -0.077 | 3.13 | 3.65% |
| B07 | 4.651 | 2.803 | 1.848 | 0.635 | 3.061 | 6.37% |
| H09 | 4.531 | 9.33 | -4.799 | -6.663 | -2.935 | 2.7% |
| L11 | 2.492 | 2.708 | -0.216 | -1.273 | 0.84 | 8.41% |
| P08 | 4.148 | 6.151 | -2.003 | -3.647 | -0.36 | 3.47% |
| S10 | 2.074 | 2.458 | -0.384 | -1.208 | 0.44 | 13.82% |
| U01 | 3.216 | 3.145 | 0.071 | -0.641 | 0.784 | 18.48% |
| U02 | 2.692 | 4.572 | -1.88 | -3.052 | -0.708 | 6.83% |
| U03 | 3.575 | 2.926 | 0.648 | -0.006 | 1.303 | 21.9% |
| **Meta analysis** | | | **0.047** | **-0.26** | **0.353** | |

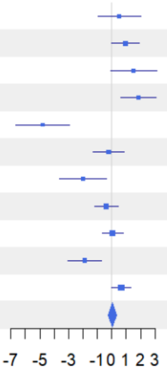


Figure S24

**Alpha with Coverage = 0.819**

| Site | PD(q=1) E | PD(q=1) C | Difference | LCL | UCL | W(fixed) |
|---|---|---|---|---|---|---|
| B04 | 3.748 | 3.164 | 0.583 | -0.227 | 1.394 | 6.56% |
| B05 | 3.022 | 2.382 | 0.641 | 0.035 | 1.247 | 11.74% |
| B06 | 3.782 | 2.658 | 1.124 | 0.21 | 2.037 | 5.16% |
| B07 | 4.064 | 2.455 | 1.609 | 0.805 | 2.412 | 6.68% |
| H09 | 3.718 | 5.193 | -1.475 | -2.442 | -0.508 | 4.61% |
| L11 | 2.401 | 2.572 | -0.171 | -0.927 | 0.585 | 7.54% |
| P08 | 3.554 | 5.262 | -1.707 | -2.921 | -0.494 | 2.93% |
| S10 | 1.985 | 2.085 | -0.1 | -0.745 | 0.545 | 10.35% |
| U01 | 2.893 | 2.85 | 0.044 | -0.453 | 0.54 | 17.47% |
| U02 | 2.485 | 4.14 | -1.655 | -2.392 | -0.918 | 7.94% |
| U03 | 3.206 | 2.725 | 0.48 | 0.004 | 0.956 | 19.02% |
| **Meta analysis** | | | **0.105** | **-0.102** | **0.313** | |

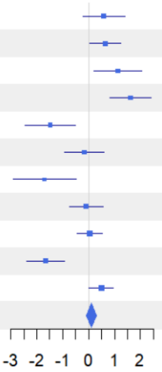


Figure S25

**Alpha with Coverage = 0.819**

| Site | PD(q=2) E | PD(q=2) C | Difference | LCL | UCL | W(fixed) |
|---|---|---|---|---|---|---|
| B04 | 3.317 | 2.631 | 0.686 | 0.153 | 1.218 | 8.21% |
| B05 | 2.631 | 2.215 | 0.417 | -0.018 | 0.851 | 12.34% |
| B06 | 3.133 | 2.283 | 0.85 | 0.284 | 1.416 | 7.28% |
| B07 | 3.586 | 2.167 | 1.419 | 0.878 | 1.96 | 7.95% |
| H09 | 2.887 | 2.825 | 0.061 | -0.555 | 0.678 | 6.13% |
| L11 | 2.287 | 2.45 | -0.163 | -0.7 | 0.374 | 8.07% |
| P08 | 2.964 | 4.442 | -1.478 | -2.407 | -0.549 | 2.7% |
| S10 | 1.899 | 1.755 | 0.144 | -0.411 | 0.699 | 7.56% |
| U01 | 2.627 | 2.64 | -0.013 | -0.422 | 0.396 | 13.92% |
| U02 | 2.216 | 3.672 | -1.456 | -1.937 | -0.975 | 10.06% |
| U03 | 2.894 | 2.571 | 0.324 | -0.06 | 0.708 | 15.79% |
| **Meta analysis** | | | **0.147** | **-0.006** | **0.299** | |

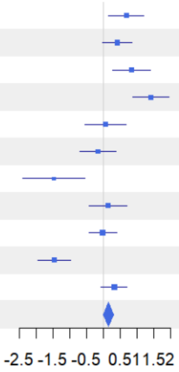


Figure S26

**Alpha with Coverage = 0.762**

| Site | FD(q=0) E | FD(q=0) C | Difference | LCL | UCL | W(fixed) |
|---|---|---|---|---|---|---|
| B04 | 3.663 | 3.334 | 0.33 | -0.715 | 1.374 | 10.75% |
| B05 | 2.886 | 2.068 | 0.818 | -0.289 | 1.925 | 9.56% |
| B06 | 3.963 | 2.804 | 1.159 | -0.081 | 2.398 | 7.63% |
| B07 | 3.63 | 2.324 | 1.306 | 0.081 | 2.532 | 7.81% |
| H09 | 3.631 | 9.679 | -6.048 | -8.372 | -3.724 | 2.17% |
| L11 | 2.687 | 2.996 | -0.309 | -1.604 | 0.985 | 7% |
| P08 | 3.562 | 5.528 | -1.966 | -3.851 | -0.081 | 3.3% |
| S10 | 2.16 | 1.923 | 0.237 | -0.798 | 1.273 | 10.93% |
| U01 | 2.962 | 2.431 | 0.531 | -0.323 | 1.385 | 16.08% |
| U02 | 3.058 | 3.695 | -0.637 | -2.129 | 0.855 | 5.27% |
| U03 | 3.274 | 2.58 | 0.694 | -0.081 | 1.47 | 19.5% |
| **Meta analysis** | | | **0.299** | **-0.043** | **0.642** | |

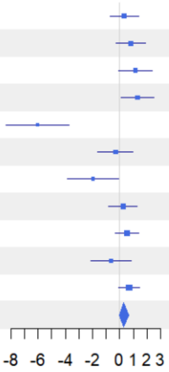


Figure S27

**Alpha with Coverage = 0.762**

| Site | FD(q=1) E | FD(q=1) C | Difference | LCL | UCL | W(fixed) |
|---|---|---|---|---|---|---|
| B04 | 3.151 | 2.65 | 0.501 | -0.266 | 1.267 | 11.17% |
| B05 | 2.395 | 1.78 | 0.615 | -0.073 | 1.303 | 13.86% |
| B06 | 3.057 | 2.252 | 0.805 | 0.02 | 1.59 | 10.65% |
| B07 | 3.168 | 1.986 | 1.182 | 0.362 | 2.002 | 9.76% |
| H09 | 3.041 | 5.904 | -2.862 | -4.306 | -1.419 | 3.15% |
| L11 | 2.343 | 2.649 | -0.306 | -1.669 | 1.057 | 3.53% |
| P08 | 3.007 | 4.832 | -1.825 | -3.381 | -0.268 | 2.71% |
| S10 | 2.017 | 1.804 | 0.214 | -0.775 | 1.202 | 6.72% |
| U01 | 2.537 | 2.132 | 0.405 | -0.24 | 1.05 | 15.76% |
| U02 | 2.378 | 3.199 | -0.821 | -1.776 | 0.134 | 7.2% |
| U03 | 2.771 | 2.208 | 0.563 | -0.088 | 1.215 | 15.47% |
| **Meta analysis** | | | **0.298** | **0.042** | **0.554** | |

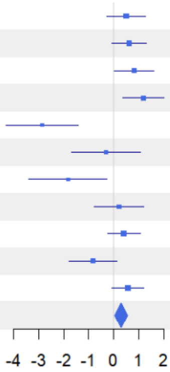


Figure S28

**Alpha with Coverage = 0.762**

| Site | FD(q=2) E | FD(q=2) C | Difference | LCL | UCL | W(fixed) |
|---|---|---|---|---|---|---|
| B04 | 2.796 | 2.234 | 0.562 | -0.05 | 1.175 | 9.29% |
| B05 | 2.086 | 1.592 | 0.494 | 0.02 | 0.968 | 15.48% |
| B06 | 2.504 | 1.874 | 0.63 | -0.055 | 1.316 | 7.42% |
| B07 | 2.813 | 1.711 | 1.102 | 0.543 | 1.661 | 11.14% |
| H09 | 2.596 | 3.072 | -0.477 | -1.415 | 0.462 | 3.95% |
| L11 | 2.074 | 2.386 | -0.312 | -1.254 | 0.631 | 3.92% |
| P08 | 2.415 | 3.899 | -1.485 | -2.541 | -0.429 | 3.12% |
| S10 | 1.866 | 1.613 | 0.253 | -0.456 | 0.962 | 6.93% |
| U01 | 2.241 | 1.924 | 0.316 | -0.128 | 0.761 | 17.62% |
| U02 | 1.879 | 2.809 | -0.93 | -1.598 | -0.261 | 7.8% |
| U03 | 2.414 | 1.956 | 0.458 | -0.053 | 0.969 | 13.33% |
| **Meta analysis** | | | **0.283** | **0.096** | **0.469** | |

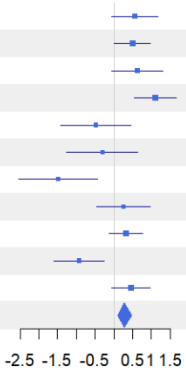


Figure S29

**Figures S3-S29** Forest plots showing meta-analysis results for bats, standardized to the 25th quantile of sample coverage values among samples extrapolated to twice their original size (0.819 for TD and PD; 0.762 for FD). Three separate meta-analyses were conducted for gamma (γ-), beta (1–S, β-), and alpha (α-) diversity, each assessed across three diversity facets: taxonomic (TD), phylogenetic (PD), and functional (FD) diversity, and three orders of q, emphasizing infrequent ($q = 0$), frequent ($q = 1$), and highly frequent ($q = 2$) species within assemblages. Site IDs refer to the 11 experimental landscapes: Lübeck (L11), Saarland (S10), University Forest (U01–U03), Passau (P08), Hunsrück-Hochwald National Park (H09), and Bavarian Forest National Park (B04–B07). E (Enhanced) and C (Control) are the estimated means of diversity for treatment and control districts. Standardized diversity differences are calculated as E - C. LCL and UCL indicate lower and upper confidence limits. Blue squares represent the standardized difference, with horizontal lines showing 95% confidence intervals (CIs). The blue diamond denotes the overall meta-analytic estimate across all landscapes. A positive value indicates higher diversity in treatment than control districts. Significance is inferred when the CI does not include zero. W(fixed) indicates the fixed-effect weight assigned to each site in the meta-analysis. All metrics are expressed in the same units of species, lineage, or functional group equivalents and can be directly compared within each diversity level (α, β, γ) (Chao et al. 2021). To account for variation in patch numbers between sites, we applied a Jaccard-type turnover transformation (1-S) of multiplicative β-diversity, which quantifies dissimilarity between assemblages relative to γ (see methods). Across levels, only α and γ can be directly and meaningfully compared.

Preprint not peer reviewed

**Gamma with Coverage = 0.991**

| Site | TD(q=0) E | TD(q=0) C | Difference | LCL | UCL | W(fixed) |
|---|---|---|---|---|---|---|
| B04 | 44.41 | 46.619 | -2.209 | -23.49 | 19.071 | 0.28% |
| B05 | 46.066 | 72.429 | -26.363 | -91.773 | 39.047 | 0.03% |
| B06 | 49.616 | 51.154 | -1.538 | -7.486 | 4.41 | 3.58% |
| B07 | 51.732 | 46.653 | 5.079 | 0.522 | 9.636 | 6.09% |
| H09 | 42.813 | 42.688 | 0.125 | -2.973 | 3.224 | 13.18% |
| L11 | 47.614 | 49.286 | -1.672 | -16.703 | 13.359 | 0.56% |
| P08 | 45.885 | 42.761 | 3.124 | 0.309 | 5.94 | 15.97% |
| S10 | 44.535 | 42.645 | 1.89 | -0.962 | 4.742 | 15.56% |
| U01 | 48.888 | 46.119 | 2.77 | -0.395 | 5.934 | 12.64% |
| U02 | 48.706 | 45.978 | 2.728 | -0.332 | 5.787 | 13.52% |
| U03 | 52.806 | 51.538 | 1.268 | -1.341 | 3.876 | 18.6% |
| **Meta analysis** | | | **1.995** | **0.87** | **3.12** | |

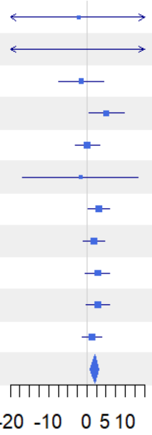


Figure S30

**Gamma with Coverage = 0.991**

| Site | TD(q=1) E | TD(q=1) C | Difference | LCL | UCL | W(fixed) |
|---|---|---|---|---|---|---|
| B04 | 31.013 | 28.006 | 3.007 | 1.448 | 4.566 | 5.09% |
| B05 | 31.845 | 31.297 | 0.548 | -1.816 | 2.911 | 2.21% |
| B06 | 37.081 | 34.523 | 2.558 | 1.302 | 3.814 | 7.84% |
| B07 | 36.33 | 30.522 | 5.808 | 4.091 | 7.525 | 4.2% |
| H09 | 33.807 | 36.189 | -2.383 | -3.331 | -1.434 | 13.76% |
| L11 | 38.527 | 37.405 | 1.121 | -0.385 | 2.627 | 5.45% |
| P08 | 36.615 | 35.546 | 1.069 | 0.087 | 2.05 | 12.84% |
| S10 | 34.916 | 31.791 | 3.125 | 2.027 | 4.222 | 10.27% |
| U01 | 40.45 | 40.734 | -0.284 | -1.277 | 0.709 | 12.56% |
| U02 | 41.337 | 38.01 | 3.327 | 2.402 | 4.252 | 14.45% |
| U03 | 44.274 | 42.831 | 1.444 | 0.399 | 2.489 | 11.32% |
| **Meta analysis** | | | **1.409** | **1.058** | **1.761** | |

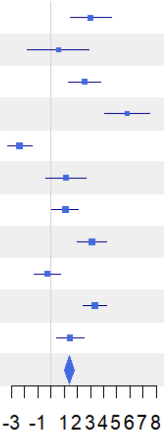


Figure S31

**Gamma with Coverage = 0.991**

| Site | TD(q=2) E | TD(q=2) C | Difference | LCL | UCL | W(fixed) |
|---|---|---|---|---|---|---|
| B04 | 27.541 | 24.164 | 3.377 | 2.433 | 4.321 | 8% |
| B05 | 27.605 | 25.521 | 2.084 | 0.8 | 3.369 | 4.32% |
| B06 | 33.786 | 30.456 | 3.331 | 2.31 | 4.351 | 6.85% |
| B07 | 32.588 | 25.929 | 6.659 | 5.418 | 7.9 | 4.63% |
| H09 | 31.062 | 34.058 | -2.996 | -3.853 | -2.139 | 9.7% |
| L11 | 36.014 | 34.705 | 1.308 | 0.474 | 2.142 | 10.25% |
| P08 | 34.124 | 33.215 | 0.909 | 0.184 | 1.634 | 13.57% |
| S10 | 32.201 | 28.681 | 3.519 | 2.701 | 4.337 | 10.66% |
| U01 | 37.801 | 38.772 | -0.971 | -1.85 | -0.093 | 9.23% |
| U02 | 39.092 | 35.223 | 3.869 | 3.162 | 4.576 | 14.25% |
| U03 | 41.205 | 40.038 | 1.166 | 0.254 | 2.079 | 8.56% |
| **Meta analysis** | | | **1.8** | **1.533** | **2.066** | |

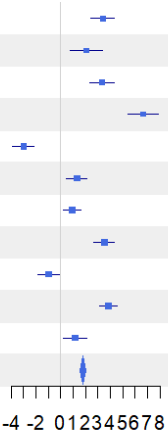

Figure S32

**Gamma with Coverage = 0.991**

| Site | PD(q=0) E | PD(q=0) C | Difference | LCL | UCL | W(fixed) |
|---|---|---|---|---|---|---|
| B04 | 14.851 | 15.307 | -0.456 | -6.93 | 6.019 | 2.12% |
| B05 | 15.324 | 22.238 | -6.914 | -12.058 | -1.77 | 3.37% |
| B06 | 16.231 | 16.514 | -0.283 | -2.685 | 2.12 | 15.43% |
| B07 | 16.933 | 16.497 | 0.435 | -5.448 | 6.319 | 2.57% |
| H09 | 13.821 | 14.066 | -0.246 | -3.092 | 2.6 | 10.99% |
| L11 | 16.68 | 16.283 | 0.397 | -5.714 | 6.509 | 2.38% |
| P08 | 15.078 | 14.521 | 0.558 | -3.295 | 4.411 | 6% |
| S10 | 14.815 | 13.863 | 0.952 | -3.836 | 5.74 | 3.88% |
| U01 | 15.609 | 14.964 | 0.645 | -1.355 | 2.646 | 22.25% |
| U02 | 15.524 | 14.977 | 0.546 | -2.162 | 3.255 | 12.14% |
| U03 | 16.541 | 16.238 | 0.303 | -1.869 | 2.475 | 18.87% |
| **Meta analysis** | | | **0.045** | **-0.898** | **0.989** | |

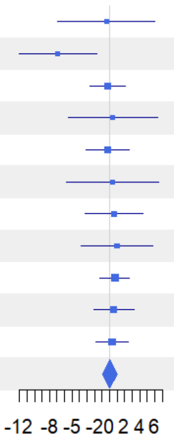


Figure S33

**Gamma with Coverage = 0.991**

| Site | PD(q=1) E | PD(q=1) C | Difference | LCL | UCL | W(fixed) |
|---|---|---|---|---|---|---|
| B04 | 11.263 | 10.388 | 0.875 | 0.285 | 1.465 | 25.56% |
| B05 | 11.511 | 11.335 | 0.175 | -2.028 | 2.378 | 1.83% |
| B06 | 13.045 | 11.957 | 1.088 | 0.12 | 2.057 | 9.47% |
| B07 | 12.568 | 11.61 | 0.957 | -3.783 | 5.698 | 0.4% |
| H09 | 11.833 | 12.402 | -0.569 | -1.132 | -0.006 | 28.05% |
| L11 | 13.875 | 13.177 | 0.698 | -4.307 | 5.703 | 0.35% |
| P08 | 12.556 | 12.657 | -0.102 | -2.945 | 2.742 | 1.1% |
| S10 | 12.232 | 11.259 | 0.973 | -2.743 | 4.689 | 0.64% |
| U01 | 13.846 | 13.928 | -0.083 | -0.664 | 0.499 | 26.25% |
| U02 | 14.016 | 13.363 | 0.653 | -1.433 | 2.74 | 2.04% |
| U03 | 14.846 | 14.466 | 0.379 | -1.06 | 1.818 | 4.29% |
| **Meta analysis** | | | **0.19** | **-0.109** | **0.488** | |

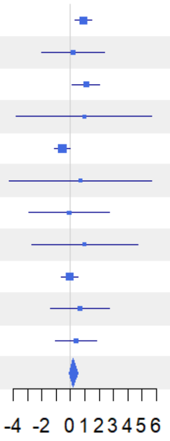


Figure S34

**Gamma with Coverage = 0.991**

| Site | PD(q=2) E | PD(q=2) C | Difference | LCL | UCL | W(fixed) |
|---|---|---|---|---|---|---|
| B04 | 10.445 | 9.419 | 1.027 | 0.7 | 1.354 | 34.32% |
| B05 | 10.411 | 9.716 | 0.695 | -1.337 | 2.727 | 0.89% |
| B06 | 12.118 | 10.953 | 1.166 | 0.305 | 2.026 | 4.96% |
| B07 | 11.587 | 10.338 | 1.249 | -3.433 | 5.931 | 0.17% |
| H09 | 11.2 | 11.871 | -0.672 | -1.027 | -0.316 | 29% |
| L11 | 13.11 | 12.482 | 0.627 | -4.448 | 5.703 | 0.14% |
| P08 | 11.855 | 11.988 | -0.133 | -2.919 | 2.652 | 0.47% |
| S10 | 11.498 | 10.481 | 1.016 | -2.612 | 4.645 | 0.28% |
| U01 | 13.322 | 13.562 | -0.24 | -0.61 | 0.13 | 26.88% |
| U02 | 13.55 | 12.799 | 0.751 | -1.313 | 2.816 | 0.86% |
| U03 | 14.264 | 13.923 | 0.341 | -1.007 | 1.689 | 2.02% |
| **Meta analysis** | | | **0.176** | **-0.016** | **0.367** | |

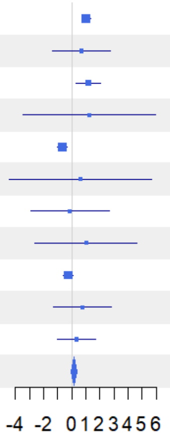

Figure S35

**Gamma with Coverage = 0.991**

| Site | FD(q=0) E | FD(q=0) C | Difference | LCL | UCL | W(fixed) |
|---|---|---|---|---|---|---|
| B04 | 25.532 | 22.034 | 3.498 | 0.38 | 6.616 | 2.06% |
| B05 | 24.524 | 24.906 | -0.382 | -3.62 | 2.856 | 1.91% |
| B06 | 30.456 | 27.439 | 3.017 | 1.555 | 4.479 | 9.35% |
| B07 | 29.759 | 23.519 | 6.241 | 4.518 | 7.963 | 6.74% |
| H09 | 28.973 | 31.509 | -2.536 | -3.951 | -1.122 | 9.99% |
| L11 | 33.81 | 32.742 | 1.068 | -1.02 | 3.155 | 4.59% |
| P08 | 33.031 | 31.111 | 1.92 | 0.808 | 3.032 | 16.16% |
| S10 | 30.26 | 25.922 | 4.338 | 2.927 | 5.748 | 10.05% |
| U01 | 35.112 | 36.162 | -1.05 | -2.66 | 0.559 | 7.72% |
| U02 | 36.728 | 32.812 | 3.916 | 2.814 | 5.018 | 16.47% |
| U03 | 38.002 | 37.033 | 0.969 | -0.186 | 2.123 | 14.99% |
| **Meta analysis** | | | **2.018** | **1.571** | **2.465** | |

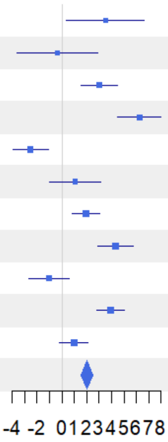


Figure S36

**Gamma with Coverage = 0.991**

| Site | FD(q=1) E | FD(q=1) C | Difference | LCL | UCL | W(fixed) |
|---|---|---|---|---|---|---|
| B04 | 23.728 | 19.667 | 4.061 | 3.351 | 4.772 | 10.97% |
| B05 | 22.975 | 20.234 | 2.741 | 1.785 | 3.696 | 6.06% |
| B06 | 28.901 | 25.22 | 3.68 | 2.881 | 4.479 | 8.67% |
| B07 | 27.673 | 21.319 | 6.354 | 5.507 | 7.202 | 7.71% |
| H09 | 27.736 | 30.701 | -2.965 | -3.763 | -2.168 | 8.71% |
| L11 | 32.764 | 31.414 | 1.35 | 0.457 | 2.243 | 6.94% |
| P08 | 31.818 | 30.104 | 1.714 | 1.058 | 2.37 | 12.88% |
| S10 | 28.96 | 24.596 | 4.364 | 3.578 | 5.15 | 8.97% |
| U01 | 34.173 | 35.482 | -1.309 | -2.046 | -0.572 | 10.2% |
| U02 | 35.851 | 31.83 | 4.021 | 3.279 | 4.764 | 10.05% |
| U03 | 37.059 | 35.964 | 1.096 | 0.305 | 1.887 | 8.85% |
| **Meta analysis** | | | **2.236** | **2.001** | **2.471** | |

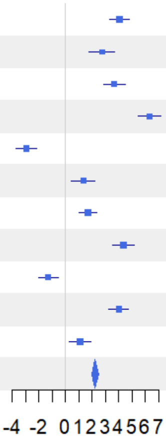


Figure S37

**Gamma with Coverage = 0.991**

| Site | FD(q=2) E | FD(q=2) C | Difference | LCL | UCL | W(fixed) |
|---|---|---|---|---|---|---|
| B04 | 23.295 | 19.172 | 4.123 | 3.506 | 4.74 | 11.92% |
| B05 | 22.512 | 19.558 | 2.954 | 2.106 | 3.801 | 6.32% |
| B06 | 28.47 | 24.697 | 3.774 | 2.961 | 4.587 | 6.86% |
| B07 | 27.176 | 20.704 | 6.471 | 5.631 | 7.311 | 6.43% |
| H09 | 27.357 | 30.438 | -3.081 | -3.716 | -2.446 | 11.25% |
| L11 | 32.481 | 31.117 | 1.364 | 0.591 | 2.136 | 7.61% |
| P08 | 31.47 | 29.763 | 1.707 | 1.137 | 2.277 | 13.96% |
| S10 | 28.609 | 24.217 | 4.391 | 3.769 | 5.013 | 11.74% |
| U01 | 33.884 | 35.243 | -1.359 | -2.037 | -0.681 | 9.87% |
| U02 | 35.576 | 31.482 | 4.094 | 3.256 | 4.933 | 6.46% |
| U03 | 36.72 | 35.615 | 1.105 | 0.33 | 1.879 | 7.57% |
| **Meta analysis** | | | **2.079** | **1.866** | **2.292** | |

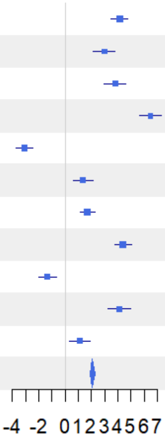

Figure S38

**1-S with Coverage = 0.991**

| Site | TD(q=0) E | TD(q=0) C | Difference | LCL | UCL | W(fixed) |
|---|---|---|---|---|---|---|
| B04 | 0.21 | 0.34 | -0.131 | -0.256 | -0.005 | 3.5% |
| B05 | 0.208 | 0.605 | -0.397 | -0.508 | -0.286 | 4.48% |
| B06 | 0.187 | 0.239 | -0.052 | -0.148 | 0.043 | 6.11% |
| B07 | 0.166 | 0.252 | -0.085 | -0.187 | 0.017 | 5.31% |
| H09 | 0.142 | 0.133 | 0.009 | -0.097 | 0.114 | 4.94% |
| L11 | 0.215 | 0.272 | -0.057 | -0.164 | 0.05 | 4.87% |
| P08 | 0.226 | 0.203 | 0.023 | -0.05 | 0.095 | 10.54% |
| S10 | 0.215 | 0.134 | 0.081 | -0.021 | 0.183 | 5.31% |
| U01 | 0.229 | 0.179 | 0.049 | -0.008 | 0.107 | 16.81% |
| U02 | 0.202 | 0.28 | -0.078 | -0.136 | -0.021 | 16.57% |
| U03 | 0.241 | 0.265 | -0.023 | -0.074 | 0.027 | 21.58% |
| **Meta analysis** | | | **-0.036** | **-0.059** | **-0.012** | |

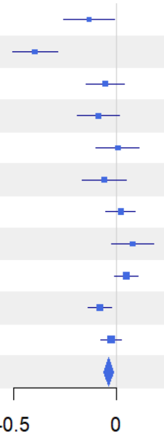


Figure S39

**1-S with Coverage = 0.991**

| Site | TD(q=1) E | TD(q=1) C | Difference | LCL | UCL | W(fixed) |
|---|---|---|---|---|---|---|
| B04 | 0.411 | 0.488 | -0.077 | -0.104 | -0.05 | 5.53% |
| B05 | 0.436 | 0.514 | -0.078 | -0.108 | -0.049 | 4.59% |
| B06 | 0.38 | 0.468 | -0.088 | -0.109 | -0.066 | 8.6% |
| B07 | 0.4 | 0.469 | -0.069 | -0.093 | -0.045 | 6.97% |
| H09 | 0.342 | 0.321 | 0.021 | 0 | 0.042 | 8.85% |
| L11 | 0.308 | 0.326 | -0.018 | -0.042 | 0.005 | 7.02% |
| P08 | 0.316 | 0.373 | -0.058 | -0.079 | -0.037 | 9.34% |
| S10 | 0.337 | 0.304 | 0.033 | 0.009 | 0.056 | 7.35% |
| U01 | 0.403 | 0.379 | 0.024 | 0.007 | 0.041 | 14.4% |
| U02 | 0.367 | 0.437 | -0.07 | -0.088 | -0.052 | 12.6% |
| U03 | 0.417 | 0.436 | -0.019 | -0.035 | -0.002 | 14.76% |
| **Meta analysis** | | | **-0.031** | **-0.037** | **-0.024** | |

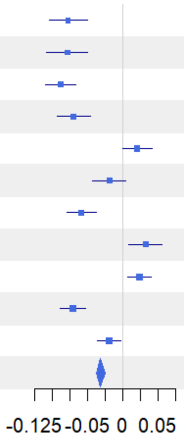


Figure S40

**1-S with Coverage = 0.991**

| Site | TD(q=2) E | TD(q=2) C | Difference | LCL | UCL | W(fixed) |
|---|---|---|---|---|---|---|
| B04 | 0.469 | 0.568 | -0.099 | -0.119 | -0.08 | 6.66% |
| B05 | 0.503 | 0.578 | -0.074 | -0.101 | -0.047 | 3.55% |
| B06 | 0.472 | 0.544 | -0.072 | -0.088 | -0.055 | 8.94% |
| B07 | 0.509 | 0.532 | -0.023 | -0.046 | 0 | 4.89% |
| H09 | 0.417 | 0.41 | 0.008 | -0.009 | 0.024 | 9.39% |
| L11 | 0.362 | 0.382 | -0.02 | -0.036 | -0.004 | 10.47% |
| P08 | 0.361 | 0.441 | -0.08 | -0.095 | -0.066 | 12.94% |
| S10 | 0.401 | 0.347 | 0.054 | 0.031 | 0.076 | 5.08% |
| U01 | 0.466 | 0.458 | 0.008 | -0.006 | 0.022 | 13.01% |
| U02 | 0.435 | 0.49 | -0.055 | -0.069 | -0.041 | 13.64% |
| U03 | 0.475 | 0.495 | -0.02 | -0.035 | -0.005 | 11.42% |
| **Meta analysis** | | | **-0.035** | **-0.04** | **-0.03** | |

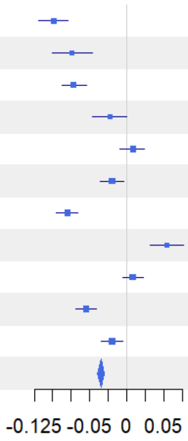

Figure S41

**1-S with Coverage = 0.991**

| Site | PD(q=0) E | PD(q=0) C | Difference | LCL | UCL | W(fixed) |
|---|---|---|---|---|---|---|
| B04 | 0.171 | 0.25 | -0.078 | -0.232 | 0.075 | 5.23% |
| B05 | 0.163 | 0.469 | -0.307 | -0.473 | -0.14 | 4.45% |
| B06 | 0.133 | 0.182 | -0.049 | -0.147 | 0.049 | 13% |
| B07 | 0.137 | 0.22 | -0.083 | -0.337 | 0.171 | 1.92% |
| H09 | 0.09 | 0.103 | -0.012 | -0.113 | 0.088 | 12.32% |
| L11 | 0.183 | 0.219 | -0.035 | -0.306 | 0.236 | 1.68% |
| P08 | 0.179 | 0.144 | 0.035 | -0.174 | 0.245 | 2.82% |
| S10 | 0.164 | 0.086 | 0.079 | -0.138 | 0.296 | 2.62% |
| U01 | 0.141 | 0.103 | 0.038 | -0.032 | 0.107 | 25.69% |
| U02 | 0.123 | 0.195 | -0.072 | -0.173 | 0.029 | 12.07% |
| U03 | 0.15 | 0.182 | -0.031 | -0.114 | 0.051 | 18.19% |
| **Meta analysis** | | | **-0.029** | **-0.065** | **0.006** | |

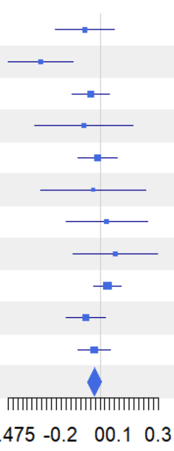


Figure 42

**1-S with Coverage = 0.991**

| Site | PD(q=1) E | PD(q=1) C | Difference | LCL | UCL | W(fixed) |
|---|---|---|---|---|---|---|
| B04 | 0.333 | 0.434 | -0.101 | -0.127 | -0.075 | 25.92% |
| B05 | 0.372 | 0.459 | -0.087 | -0.25 | 0.077 | 0.65% |
| B06 | 0.341 | 0.395 | -0.053 | -0.099 | -0.008 | 8.32% |
| B07 | 0.353 | 0.426 | -0.073 | -0.449 | 0.304 | 0.12% |
| H09 | 0.278 | 0.262 | 0.015 | -0.009 | 0.04 | 28.73% |
| L11 | 0.285 | 0.281 | 0.004 | -0.381 | 0.389 | 0.12% |
| P08 | 0.264 | 0.323 | -0.058 | -0.283 | 0.166 | 0.35% |
| S10 | 0.286 | 0.258 | 0.029 | -0.291 | 0.348 | 0.17% |
| U01 | 0.31 | 0.296 | 0.014 | -0.01 | 0.037 | 31.69% |
| U02 | 0.286 | 0.367 | -0.081 | -0.213 | 0.051 | 1% |
| U03 | 0.321 | 0.343 | -0.023 | -0.1 | 0.055 | 2.93% |
| **Meta analysis** | | | **-0.024** | **-0.037** | **-0.011** | |

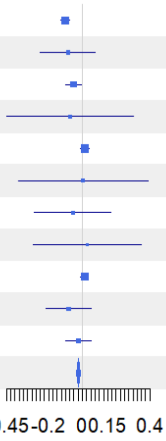


Figure S43

**1-S with Coverage = 0.991**

| Site | PD(q=2) E | PD(q=2) C | Difference | LCL | UCL | W(fixed) |
|---|---|---|---|---|---|---|
| B04 | 0.398 | 0.527 | -0.128 | -0.145 | -0.111 | 30.54% |
| B05 | 0.433 | 0.52 | -0.087 | -0.269 | 0.095 | 0.26% |
| B06 | 0.413 | 0.468 | -0.055 | -0.098 | -0.012 | 4.69% |
| B07 | 0.448 | 0.498 | -0.049 | -0.491 | 0.392 | 0.05% |
| H09 | 0.343 | 0.329 | 0.014 | -0.003 | 0.031 | 30.52% |
| L11 | 0.329 | 0.327 | 0.001 | -0.426 | 0.428 | 0.05% |
| P08 | 0.295 | 0.373 | -0.078 | -0.312 | 0.156 | 0.16% |
| S10 | 0.338 | 0.304 | 0.035 | -0.31 | 0.38 | 0.07% |
| U01 | 0.367 | 0.373 | -0.006 | -0.023 | 0.01 | 31.72% |
| U02 | 0.348 | 0.431 | -0.083 | -0.219 | 0.054 | 0.47% |
| U03 | 0.378 | 0.396 | -0.018 | -0.096 | 0.059 | 1.47% |
| **Meta analysis** | | | **-0.04** | **-0.05** | **-0.031** | |

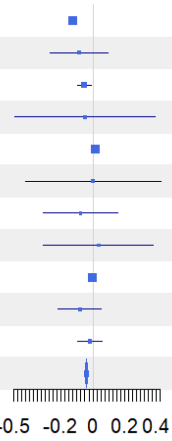

Figure S44

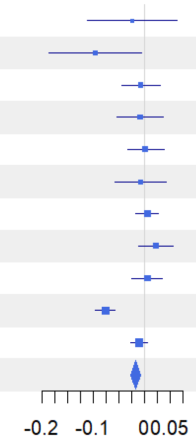


| Site | FD(q=0) E | FD(q=0) C | Difference | LCL | UCL | W(fixed) |
|---|---|---|---|---|---|---|
| B04 | 0.026 | 0.051 | -0.025 | -0.112 | 0.062 | 1.02% |
| B05 | 0.024 | 0.121 | -0.097 | -0.188 | -0.006 | 0.92% |
| B06 | 0.024 | 0.031 | -0.008 | -0.045 | 0.03 | 5.47% |
| B07 | 0.02 | 0.029 | -0.009 | -0.054 | 0.036 | 3.81% |
| H09 | 0.017 | 0.016 | 0.001 | -0.034 | 0.037 | 6.15% |
| L11 | 0.027 | 0.035 | -0.008 | -0.059 | 0.042 | 3.02% |
| P08 | 0.032 | 0.028 | 0.005 | -0.018 | 0.027 | 15.41% |
| S10 | 0.031 | 0.01 | 0.021 | -0.013 | 0.055 | 6.58% |
| U01 | 0.026 | 0.021 | 0.005 | -0.026 | 0.035 | 8.34% |
| U02 | 0.025 | 0.103 | -0.077 | -0.097 | -0.058 | 19.83% |
| U03 | 0.027 | 0.038 | -0.011 | -0.027 | 0.005 | 29.44% |
| **Meta analysis** | | | **-0.018** | **-0.027** | **-0.009** | |

Figure S45

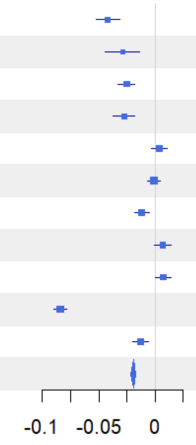


| Site | FD(q=1) E | FD(q=1) C | Difference | LCL | UCL | W(fixed) |
|---|---|---|---|---|---|---|
| B04 | 0.085 | 0.127 | -0.042 | -0.052 | -0.031 | 4.38% |
| B05 | 0.098 | 0.128 | -0.029 | -0.044 | -0.014 | 2.17% |
| B06 | 0.085 | 0.11 | -0.025 | -0.033 | -0.018 | 8.84% |
| B07 | 0.094 | 0.121 | -0.027 | -0.037 | -0.018 | 5.38% |
| H09 | 0.073 | 0.069 | 0.004 | -0.003 | 0.011 | 10.12% |
| L11 | 0.054 | 0.055 | -0.001 | -0.007 | 0.005 | 12.85% |
| P08 | 0.061 | 0.072 | -0.012 | -0.018 | -0.005 | 13.7% |
| S10 | 0.063 | 0.056 | 0.007 | -0.001 | 0.014 | 8.93% |
| U01 | 0.075 | 0.068 | 0.007 | 0 | 0.014 | 9.56% |
| U02 | 0.067 | 0.151 | -0.084 | -0.09 | -0.078 | 14.06% |
| U03 | 0.08 | 0.093 | -0.013 | -0.02 | -0.006 | 10.01% |
| **Meta analysis** | | | **-0.019** | **-0.022** | **-0.017** | |

Figure S46

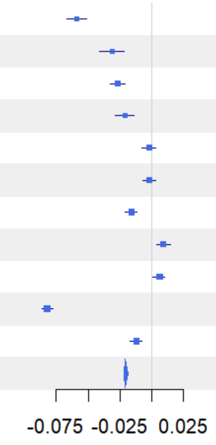


| Site | FD(q=2) E | FD(q=2) C | Difference | LCL | UCL | W(fixed) |
|---|---|---|---|---|---|---|
| B04 | 0.104 | 0.162 | -0.059 | -0.067 | -0.051 | 3.86% |
| B05 | 0.126 | 0.157 | -0.031 | -0.041 | -0.022 | 2.86% |
| B06 | 0.11 | 0.137 | -0.027 | -0.033 | -0.021 | 6.51% |
| B07 | 0.125 | 0.147 | -0.021 | -0.029 | -0.014 | 4.14% |
| H09 | 0.092 | 0.095 | -0.002 | -0.008 | 0.003 | 9.51% |
| L11 | 0.069 | 0.071 | -0.002 | -0.007 | 0.003 | 10.8% |
| P08 | 0.072 | 0.088 | -0.016 | -0.021 | -0.012 | 13.38% |
| S10 | 0.078 | 0.069 | 0.009 | 0.004 | 0.015 | 8.75% |
| U01 | 0.095 | 0.09 | 0.006 | 0.001 | 0.01 | 14.34% |
| U02 | 0.085 | 0.167 | -0.082 | -0.086 | -0.078 | 15.22% |
| U03 | 0.101 | 0.113 | -0.012 | -0.017 | -0.008 | 10.62% |
| **Meta analysis** | | | **-0.021** | **-0.022** | **-0.019** | |

Figure S47

**Alpha with Coverage = 0.991**

| Site | TD(q=0) E | TD(q=0) C | Difference | LCL | UCL | W(fixed) |
|---|---|---|---|---|---|---|
| B04 | 36.128 | 32.514 | 3.614 | 0.298 | 6.929 | 3.64% |
| B05 | 37.536 | 34.084 | 3.452 | -1.402 | 8.306 | 1.7% |
| B06 | 41.499 | 40.267 | 1.232 | -2.046 | 4.511 | 3.73% |
| B07 | 44.199 | 36.222 | 7.977 | 2.543 | 13.412 | 1.36% |
| H09 | 37.411 | 37.625 | -0.214 | -2.456 | 2.028 | 7.97% |
| L11 | 38.521 | 37.377 | 1.144 | -0.982 | 3.269 | 8.87% |
| P08 | 36.676 | 35.035 | 1.641 | -0.048 | 3.33 | 14.04% |
| S10 | 36.023 | 38.072 | -2.049 | -6.362 | 2.265 | 2.15% |
| U01 | 38.455 | 38.395 | 0.06 | -1.433 | 1.552 | 17.98% |
| U02 | 39.531 | 34.012 | 5.519 | 4.213 | 6.825 | 23.49% |
| U03 | 40.911 | 38.803 | 2.108 | 0.478 | 3.739 | 15.07% |
| **Meta analysis** | | | **2.24** | **1.607** | **2.873** | |

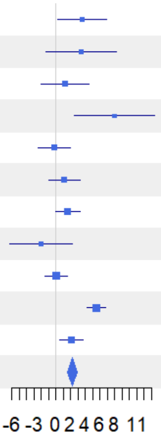


Figure S48

**Alpha with Coverage = 0.991**

| Site | TD(q=1) E | TD(q=1) C | Difference | LCL | UCL | W(fixed) |
|---|---|---|---|---|---|---|
| B04 | 19.681 | 15.856 | 3.825 | 3.335 | 4.315 | 9.34% |
| B05 | 19.509 | 17.223 | 2.287 | 1.584 | 2.989 | 4.54% |
| B06 | 24.743 | 20.162 | 4.581 | 3.995 | 5.167 | 6.52% |
| B07 | 23.618 | 17.79 | 5.828 | 5.059 | 6.596 | 3.8% |
| H09 | 23.516 | 25.848 | -2.332 | -2.894 | -1.77 | 7.1% |
| L11 | 27.989 | 26.559 | 1.43 | 0.969 | 1.89 | 10.58% |
| P08 | 26.342 | 23.747 | 2.595 | 2.144 | 3.045 | 11.04% |
| S10 | 24.464 | 24.056 | 0.408 | -0.139 | 0.956 | 7.48% |
| U01 | 25.234 | 26.325 | -1.091 | -1.503 | -0.678 | 13.17% |
| U02 | 27.184 | 22.603 | 4.582 | 4.198 | 4.965 | 15.26% |
| U03 | 27.041 | 25.402 | 1.638 | 1.19 | 2.086 | 11.17% |
| **Meta analysis** | | | **2.022** | **1.873** | **2.172** | |

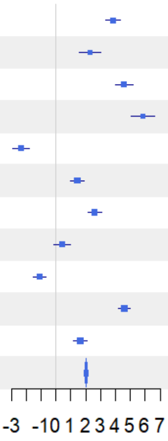


Figure S49

**Alpha with Coverage = 0.991**

| Site | TD(q=2) E | TD(q=2) C | Difference | LCL | UCL | W(fixed) |
|---|---|---|---|---|---|---|
| B04 | 16.054 | 11.955 | 4.098 | 3.715 | 4.482 | 8.57% |
| B05 | 15.255 | 12.625 | 2.631 | 2.14 | 3.121 | 5.24% |
| B06 | 19.822 | 15.732 | 4.09 | 3.641 | 4.538 | 6.27% |
| B07 | 18.076 | 13.661 | 4.416 | 3.856 | 4.975 | 4.02% |
| H09 | 19.538 | 21.653 | -2.115 | -2.6 | -1.63 | 5.36% |
| L11 | 24.432 | 22.93 | 1.502 | 1.157 | 1.847 | 10.6% |
| P08 | 23.176 | 20.183 | 2.993 | 2.664 | 3.322 | 11.65% |
| S10 | 20.734 | 20.724 | 0.01 | -0.503 | 0.524 | 4.78% |
| U01 | 21.375 | 22.203 | -0.828 | -1.116 | -0.539 | 15.15% |
| U02 | 23.227 | 19.204 | 4.023 | 3.735 | 4.311 | 15.24% |
| U03 | 22.926 | 21.529 | 1.397 | 1.087 | 1.707 | 13.11% |
| **Meta analysis** | | | **1.989** | **1.877** | **2.102** | |

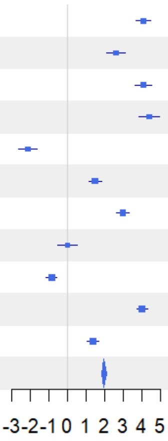

Figure S50

**Alpha with Coverage = 0.991**

| Site | PD(q=0) E | PD(q=0) C | Difference | LCL | UCL | W(fixed) |
|---|---|---|---|---|---|---|
| B04 | 12.587 | 11.906 | 0.681 | -0.522 | 1.883 | 3.63% |
| B05 | 13.109 | 13.106 | 0.003 | -1.487 | 1.492 | 2.36% |
| B06 | 14.344 | 13.844 | 0.499 | -0.672 | 1.671 | 3.82% |
| B07 | 14.898 | 13.267 | 1.63 | -0.24 | 3.5 | 1.5% |
| H09 | 12.712 | 12.782 | -0.071 | -0.856 | 0.715 | 8.5% |
| L11 | 13.962 | 13.118 | 0.844 | -0.253 | 1.94 | 4.36% |
| P08 | 12.681 | 12.665 | 0.016 | -0.607 | 0.64 | 13.5% |
| S10 | 12.651 | 12.913 | -0.263 | -1.573 | 1.047 | 3.06% |
| U01 | 13.559 | 13.527 | 0.032 | -0.481 | 0.545 | 19.94% |
| U02 | 13.736 | 12.259 | 1.477 | 0.991 | 1.964 | 22.18% |
| U03 | 14.22 | 13.487 | 0.734 | 0.181 | 1.287 | 17.16% |
| **Meta analysis** | | | **0.553** | **0.324** | **0.782** | |

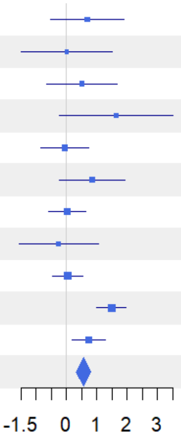


Figure S51

**Alpha with Coverage = 0.991**

| Site | PD(q=1) E | PD(q=1) C | Difference | LCL | UCL | W(fixed) |
|---|---|---|---|---|---|---|
| B04 | 7.93 | 6.378 | 1.552 | 1.361 | 1.743 | 11.27% |
| B05 | 7.702 | 6.784 | 0.918 | 0.626 | 1.21 | 4.84% |
| B06 | 9.151 | 7.763 | 1.388 | 1.127 | 1.65 | 6.03% |
| B07 | 8.682 | 7.213 | 1.469 | 0.843 | 2.095 | 1.05% |
| H09 | 8.914 | 9.51 | -0.596 | -0.766 | -0.427 | 14.41% |
| L11 | 10.358 | 9.883 | 0.475 | -0.371 | 1.32 | 0.58% |
| P08 | 9.608 | 9.028 | 0.58 | 0.296 | 0.864 | 5.12% |
| S10 | 9.118 | 8.939 | 0.179 | -0.419 | 0.777 | 1.15% |
| U01 | 9.845 | 10.083 | -0.239 | -0.368 | -0.11 | 24.89% |
| U02 | 10.273 | 8.805 | 1.468 | 1.298 | 1.638 | 14.29% |
| U03 | 10.404 | 9.832 | 0.572 | 0.413 | 0.731 | 16.37% |
| **Meta analysis** | | | **0.511** | **0.447** | **0.575** | |

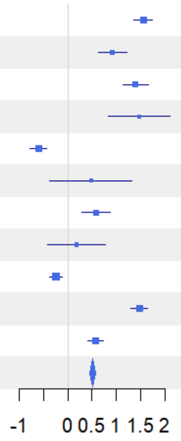


Figure S52

**Alpha with Coverage = 0.991**

| Site | PD(q=2) E | PD(q=2) C | Difference | LCL | UCL | W(fixed) |
|---|---|---|---|---|---|---|
| B04 | 6.746 | 5.009 | 1.737 | 1.6 | 1.874 | 14.81% |
| B05 | 6.4 | 5.292 | 1.108 | 0.895 | 1.321 | 6.14% |
| B06 | 7.741 | 6.4 | 1.341 | 1.117 | 1.566 | 5.52% |
| B07 | 7.042 | 5.766 | 1.276 | 0.716 | 1.835 | 0.89% |
| H09 | 7.785 | 8.4 | -0.615 | -0.751 | -0.479 | 15.07% |
| L11 | 9.281 | 8.852 | 0.429 | -0.39 | 1.248 | 0.41% |
| P08 | 8.751 | 8.015 | 0.735 | 0.464 | 1.007 | 3.77% |
| S10 | 8.039 | 7.935 | 0.104 | -0.46 | 0.667 | 0.88% |
| U01 | 8.762 | 8.841 | -0.079 | -0.187 | 0.028 | 24.11% |
| U02 | 9.151 | 7.682 | 1.469 | 1.316 | 1.623 | 11.83% |
| U03 | 9.233 | 8.776 | 0.458 | 0.328 | 0.587 | 16.58% |
| **Meta analysis** | | | **0.579** | **0.526** | **0.632** | |

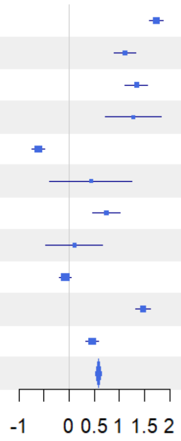

Figure S53

**Alpha with Coverage = 0.991**

| Site | FD(q=0) E | FD(q=0) C | Difference | LCL | UCL | W(fixed) |
|---|---|---|---|---|---|---|
| B04 | 24.737 | 20.626 | 4.111 | 3.104 | 5.119 | 6.89% |
| B05 | 23.792 | 21.234 | 2.558 | 1.194 | 3.922 | 3.76% |
| B06 | 29.64 | 26.369 | 3.271 | 2.249 | 4.293 | 6.7% |
| B07 | 29.033 | 22.591 | 6.441 | 5.12 | 7.763 | 4.01% |
| H09 | 28.406 | 30.977 | -2.571 | -3.428 | -1.714 | 9.53% |
| L11 | 32.903 | 31.584 | 1.319 | 0.508 | 2.129 | 10.65% |
| P08 | 31.972 | 30.243 | 1.729 | 1.066 | 2.392 | 15.91% |
| S10 | 29.278 | 25.613 | 3.665 | 2.652 | 4.679 | 6.81% |
| U01 | 34.178 | 35.4 | -1.222 | -1.966 | -0.479 | 12.66% |
| U02 | 35.792 | 29.589 | 6.203 | 5.488 | 6.918 | 13.68% |
| U03 | 36.945 | 35.626 | 1.319 | 0.456 | 2.181 | 9.41% |
| **Meta analysis** | | | **2.095** | **1.83** | **2.359** | |

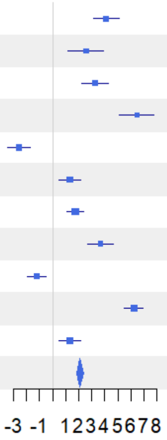


Figure S54

**Alpha with Coverage = 0.991**

| Site | FD(q=1) E | FD(q=1) C | Difference | LCL | UCL | W(fixed) |
|---|---|---|---|---|---|---|
| B04 | 22.056 | 17.636 | 4.419 | 3.839 | 5 | 11.25% |
| B05 | 21.11 | 18.09 | 3.019 | 2.283 | 3.755 | 7% |
| B06 | 26.922 | 22.963 | 3.959 | 3.27 | 4.648 | 7.99% |
| B07 | 25.553 | 19.177 | 6.376 | 5.654 | 7.098 | 7.27% |
| H09 | 26.074 | 28.969 | -2.895 | -3.577 | -2.213 | 8.16% |
| L11 | 31.293 | 29.998 | 1.295 | 0.541 | 2.049 | 6.67% |
| P08 | 30.25 | 28.373 | 1.877 | 1.339 | 2.415 | 13.12% |
| S10 | 27.445 | 23.492 | 3.953 | 3.273 | 4.632 | 8.22% |
| U01 | 32.102 | 33.513 | -1.411 | -2.02 | -0.803 | 10.25% |
| U02 | 33.873 | 27.602 | 6.27 | 5.671 | 6.87 | 10.57% |
| U03 | 34.659 | 33.264 | 1.395 | 0.762 | 2.027 | 9.49% |
| **Meta analysis** | | | **2.56** | **2.365** | **2.755** | |

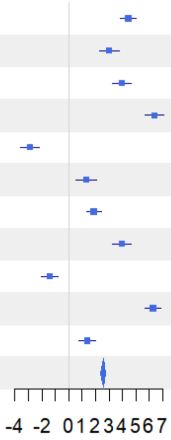


Figure S55

**Alpha with Coverage = 0.991**

| Site | FD(q=2) E | FD(q=2) C | Difference | LCL | UCL | W(fixed) |
|---|---|---|---|---|---|---|
| B04 | 21.479 | 16.94 | 4.539 | 4.016 | 5.062 | 12.12% |
| B05 | 20.429 | 17.328 | 3.101 | 2.373 | 3.828 | 6.25% |
| B06 | 26.179 | 22.222 | 3.957 | 3.264 | 4.649 | 6.9% |
| B07 | 24.701 | 18.469 | 6.233 | 5.525 | 6.94 | 6.62% |
| H09 | 25.463 | 28.292 | -2.829 | -3.384 | -2.273 | 10.74% |
| L11 | 30.738 | 29.44 | 1.298 | 0.616 | 1.979 | 7.14% |
| P08 | 29.749 | 27.835 | 1.915 | 1.437 | 2.392 | 14.53% |
| S10 | 26.888 | 22.988 | 3.9 | 3.352 | 4.447 | 11.07% |
| U01 | 31.47 | 32.857 | -1.387 | -1.968 | -0.805 | 9.8% |
| U02 | 33.267 | 27.046 | 6.221 | 5.526 | 6.915 | 6.87% |
| U03 | 33.966 | 32.585 | 1.381 | 0.736 | 2.026 | 7.96% |
| **Meta analysis** | | | **2.33** | **2.148** | **2.512** | |

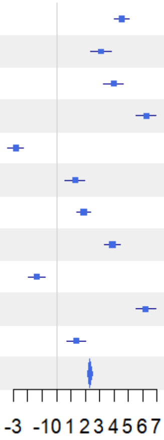

Figure S56

**Figures S30-S56** Forest plots showing meta-analysis results for birds, standardized to the minimum sample coverage across all samples when each is extrapolated to twice their original size (0.991). This level (Cmax) represents the highest coverage level at which reliable comparisons can be made across all assemblages. Three separate meta-analyses were conducted for gamma (γ-), beta (1–S, β-), and alpha (α-) diversity, each assessed across three diversity facets: taxonomic (TD), phylogenetic (PD), and functional (FD) diversity, and three orders of q, emphasizing infrequent ($q = 0$), frequent ($q = 1$), and highly frequent ($q = 2$) species within assemblages. Forest plot elements and interpretation follow the same structure described in the caption of Figures S3–S29.

Preprint not peer reviewed

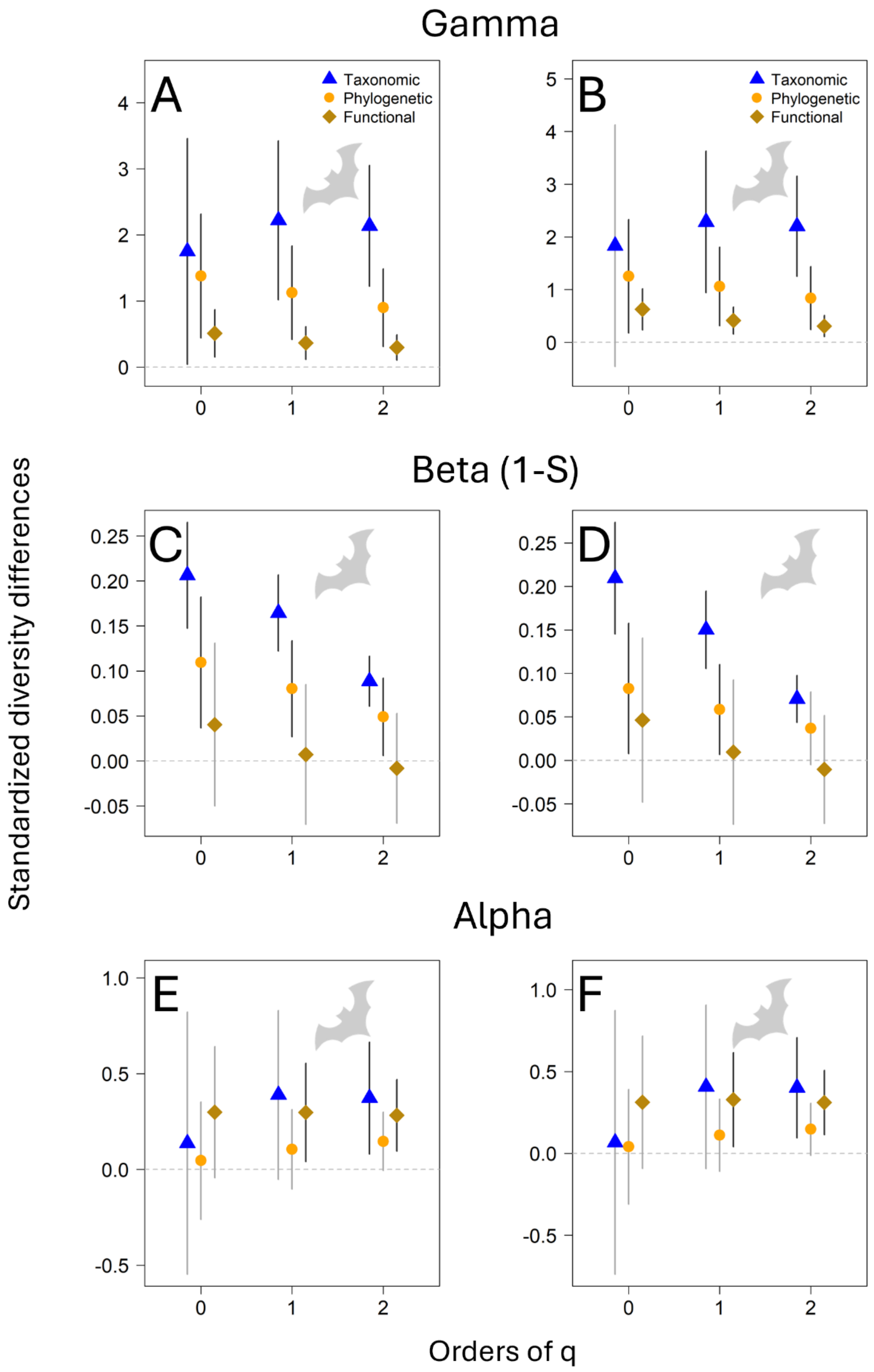


**Figure S57** Results from three meta-analyses evaluating γ-, β-, and α-diversity of bats across 11 experimental forest landscapes, using two sample coverage thresholds. Left panels show results based on the 25th quantile of sample coverage (0.819 for TD and PD; 0.762 for FD), whereas the right panels show results based on the median of sample coverage across all samples extrapolated to double size (0.888 for TD and PD; 0.836 for FD). The consistency of patterns between thresholds supports the robustness of the results and the reliability of derived inferences.

**Table S3** Overview of bat sampling units per site, treatment (E = Enhanced, C = Control), and patch. Sampling units per patch are the number of sampled nights.

| **Site** | **B04** | | **B05** | | **B06** | | **B07** | | **H09** | | **L11** | |
|---|---|---|---|---|---|---|---|---|---|---|---|---|
| **Treatment** | E | C | E | C | E | C | E | C | E | C | E | C |
| **Patches** | 9 | 9 | 9 | 9 | 9 | 9 | 9 | 8 | 9 | 9 | 9 | 9 |
| Number of sampling units in patches (patches x sampling units) | 9 × 4 | 9 × 4 | 9 × 4 | 9 × 4 | 9 × 4 | 9 × 4 | 9 × 4 | 8 × 4 | 9 × 4 | 9 × 4 | 9 × 4 | 9 × 4 |

| **Site** | **P08** | | **S10** | | **U01** | | **U02** | | **U03** | |
|---|---|---|---|---|---|---|---|---|---|---|
| **Treatments** | E | C | E | C | E | C | E | C | E | C |
| **Patches** | 8 | 9 | 9 | 8 | 15 | 15 | 14 | 14 | 15 | 15 |
| Number of sampling units in patches (patches x sampling units) | 8 × 4 | 9 × 4 | 9 × 4 | 8 × 4 | 15 × 4 | 15 × 4 | 14 × 4 | 14 × 4 | 15 × 4 | 15 × 4 |

Preprint not peer reviewed

**Table S4** Overview of bird sampling units per site, treatment (E = Enhanced, C = Control), and patch. Sampling units per patch are the number of sampled days.

| **Site** | **B04** | | **B05** | | **B06** | | **B07** | | **H09** | | **L11** | |
|---|---|---|---|---|---|---|---|---|---|---|---|---|
| **Treatment** | E | C | E | C | E | C | E | C | E | C | E | C |
| **Patches** | 9 | 9 | 9 | 8 | 8 | 9 | 8 | 9 | 9 | 9 | 9 | 9 |
| Number of sampling units in patches (patches x sampling units) | 9 × 84 | 9 × 84 | 9 × 69 | 8 × 49 | 8 × 84 | 9 × 75 | 8 × 57 | 9 × 71 | 9 × 77 | 9 × 62 | 9 × 78 | 9 × 48 |

| **Site** | **P08** | | **S10** | | **U01** | | **U02** | | **U03** | |
|---|---|---|---|---|---|---|---|---|---|---|
| **Treatments** | E | C | E | C | E | C | E | C | E | C |
| **Patches** | 9 | 9 | 9 | 5 | 15 | 15 | 15 | 14 | 15 | 15 |
| Number of sampling units in patches (patches x sampling units) | 9 × 84 | 9 × 84 | 9 × 83 | 5 × 64 | 15 × 69 | 15 × 77 | 15 × 55 | 14 × 84 | 15 × 51 | 15 × 84 |

Preprint not peer reviewed
